# Steroidal glycoalkaloids contribute to anthracnose resistance in *Solanum lycopersicum*

**DOI:** 10.1101/2022.08.08.503224

**Authors:** Matthew L. Fabian, Chong Zhang, Jianghao Sun, Neil P. Price, Pei Chen, Christopher R. Clarke, Richard W. Jones, John R. Stommel

## Abstract

Anthracnose is a widespread plant disease caused by various species of the fungal pathogen *Colletotrichum*. In solanaceous plants such as tomato (*Solanum lycoperiscum*), *Colletotrichum* infections exhibit a quiescent, asymptomatic state in developing fruit, followed by a transition to necrotrophic infections in ripe fruit. Through analysis of fruit tissue extracts of 95L368, a tomato breeding line that yields fruit with enhanced anthracnose resistance, we identified a role for steroidal glycoalkaloids (SGAs) in anthracnose resistance. The SGA α-tomatine and several of its derivatives accumulated at higher levels, in comparison to fruit of the susceptible tomato cultivar US28, 95L368 fruit extracts with fungistatic activity against *Colletotrichum*. Correspondingly, ripe and unripe 95L368 fruit displayed enhanced expression of glycoalkaloid metabolic enzyme (GAME) genes, which encode key enzymes in SGA biosynthesis. Metabolomics analysis incorporating recombinant inbred lines (RILs) generated from 95L368 and US28 yielded strong positive correlations between anthracnose resistance and accumulation of α-tomatine and several derivatives. Lastly, transient silencing of expression of the GAME genes *GAME31* and *GAME5* in anthracnose-susceptible tomato fruit yielded enhancements to anthracnose resistance. Together, our data support a role for SGAs in anthracnose defense in tomato, with a distinct SGA metabolomic profile conferring resistance to virulent *Colletotrichum* infections in ripe fruit.

**Highlight:** This work describes an important role for steroidal glycoalkaloids (SGAs), abundant secondary metabolites in tomato fruit, in defense against anthracnose, a widespread fungal disease that impacts diverse crop species.

## Introduction

Anthracnose is a global plant disease caused by *Colletotrichum* spp., hemibiotrophic fungal pathogens with a wide and diverse host range, including a variety of commercially important monocot and dicot crops such as tomato, maize, strawberry, and soybean (Dias et al., 2019; Jacobs et al., 2019; Prochno et al., 2016; Stommel and Haynes, 1998). Plant pathogens in both the field and post-harvest, *Colletotrichum* spp. encompass approximately 190 species in eleven species complexes, exhibiting multiple infection strategies in various types of plant tissue, including foliar, fruit, and root tissue, depending on the pathosystem (Baroncelli et al., 2017; Boufleur et al., 2021; Jayawardena et al., 2016). In solanaceous crops such as pepper and tomato, anthracnose is caused primarily by the species *C. coccodes, C. gloeosporioides*, *C. dematium*, and *C. acutatum*, and is characterized by the formation of necrotrophic lesions on fruit (Chapin et al., 2006; Harp et al., 2014; Pardo-De la Hoz et al., 2016). *Colletotrichum* infections of tomato fruit exhibit a transition from quiescence to virulence during fruit ripening, yielding anthracnose symptoms on ripe fruit. After overwintering in soil, sclerotia form conidia capable of germinating on immature fruit, resulting in an asymptomatic, quiescent state characterized by the formation of appressoria that are restricted to the cuticle. As infected fruit ripens, *Colletotrichum* forms hyphae that penetrate the pericarp, and produces circular depressions that progress into necrotrophic lesions on mature, ripe fruit (Alkan et al., 2015; Peres et al., 2005; Sanogo et al., 2003). In the field, traditional strategies to mitigate anthracnose infection, such as fungicide application, provide incomplete protection, thus the development of genetic tools to improve anthracnose resistance is of particular interest (Byrne et al., 1997; Wyenandt et al., 2009). Anthracnose resistance has been identified in fruits of unadapted tomato germplasm, and attempts to breed this trait into elite materials convey a polygenic, additive mode of inheritance, as well as difficulty in transfer to commercial cultivars with an intensity equal to that observed in the resistant germplasm (Stommel, 2001; Stommel and Haynes, 1998). Therefore, more targeted approaches to develop enhanced anthracnose resistance in commercial cultivars are desirable.

Steroidal glycoalkaloids (SGAs) are secondary metabolites, derived from the mevalonate biosynthetic pathway and characterized by a nitrogenous aglycone backbone and variable glycone sidechains at the C-3 position, that accumulate to high levels in solanaceous plants such as tomato, eggplant, and potato (Cárdenas et al., 2015; You and Kan, 2021; Zhao et al., 2021). Found in various plant tissues, including leaves, fruit, and tubers, some SGAs are generally classified as antinutritional compounds due to the observed toxicity in humans, when ingested at high levels, but SGAs have also been described for their potential health benefits, including anti-inflammatory, anticarcinogenic, and antioxidative properties (Friedman, 2006, 2013; Wang et al., 2018). For example, in vitro and mouse model studies have reported activation of cancer cell apoptosis and reduction of tumor cell growth following treatments of α-tomatine, the most abundant SGA in tomato leaves and green fruit (Chao et al., 2012; Kim et al., 2015). Furthermore, numerous studies have characterized α-tomatine in responses to diverse sources of biotic stress in *Solanaceae* (Hoagland, 2009; Sandrock and VanEtten, 1998). Antifungal activity of α-tomatine was observed in vitro and in tomatoes infected with *Fusarium oxysporum*, a vascular fungal pathogen and the causal agent of fusarium wilt (Ito et al., 2007; Pareja-Jaime et al., 2008), and α-tomatine was shown to restrict in vitro growth of *Phytophthora infestans*, the oomycete pathogen that causes late blight in potato (Dahlin et al., 2017; Sandrock and VanEtten, 1998). Notably, a role for α-tomatine in defense against *Colletotrichum spp*. has been described, with α-tomatine treatment observed to restrict the growth of both *C. coccodes* and *C. acutatum*, and a transgenic tomato line with attenuated levels of α-tomatine displaying enhanced susceptibility to *C. coccodes* (Itkin et al., 2011; Sandrock and VanEtten, 1998). Accordingly, SGAs, including α-tomatine, are of particular interest in the development of genetic tools to imbue enhanced levels of defense against anthracnose and other biotic diseases of tomato.

While α-tomatine accumulates to high levels in immature fruit, levels of α-tomatine and certain derivative SGAs decline together during ripening, ultimately giving way to the derivative esculeosides, end products of the SGA pathway that accumulate tFig. o high levels in ripe fruit. GLYCOALKALOID METABOLIC ENZYME (GAME) genes encode enzymes that catalyze reactions in the SGA pathway (Fig. S1) (Akiyama et al., 2019; Cárdenas et al., 2019; Lee et al., 2019; Nakayasu et al., 2020; Sonawane et al., 2018). The UDP-glycosyltransferase GAME2 catalyzes the synthesis of α-tomatine, as well as the closely related dehydrotomatine, representing a branch point for the synthesis of downstream SGA derivatives (Iijima et al., 2013; Itkin et al., 2013; Kozukue et al., 2004). As tomato fruit ripens, these compounds are converted to derivative SGAs via additional GAME proteins with hydroxylation, acetyltransferase, and glycosylation activities. For example, *GAME31* encodes a 2-oxoglutarate-dependent dioxygenase (2-ODD) that synthesizes hydroxytomatine and hydroxy-dehydrotomatine from α-tomatine and dehydrotomatine, respectively (Nakayasu et al., 2020; Szymański et al., 2020). Additional, as-yet-undiscovered GAME proteins catalyze the acetoxylation of these substrates, respectively forming the derivative acetoxytomatine and acetoxy-dehydrotomatine, and the subsequent hydroxylation reactions that result in the formation of acetoxy-hydroxytomatine and acetoxy-hydroxy-dehydrotomatine, accordingly (Cárdenas et al., 2015; Zhao et al., 2021). The uridine diphosphate (UDP) glycosyltransferase GAME5 synthesizes the formation of esculeoside A and dehydro-esculeoside A from acetoxy-hydroxytomatine and acetoxy-hydroxy-dehydrotomatine, respectively (Iijima et al., 2009; Nakayasu et al., 2020; Szymański et al., 2020). Although the ripening process in tomato is generally characterized by a decline in α-tomatine levels and coincident accumulation of the end-product esculeosides, comparisons of commercial tomato cultivars, wild tomato accessions, and derivative introgression lines have illustrated a diversity of SGA metabolomic profiles in ripe red fruit, with some lines exhibiting high levels of α-tomatine in both green and red fruit (Dzakovich et al., 2021; Iijima et al., 2013; Szymański et al., 2020; Zhu et al., 2018). While a growing body of evidence supports a role for α-tomatine in pathogen defense, little is known of the contributions of derivative SGAs, therefore germplasm of variant SGA metabolomic profiles can be particularly useful in further research to elucidate the role of SGAs in disease resistance in tomato.

Here our work incorporates an anthracnose-susceptible tomato cultivar (US28) and a small-fruited, commercially unadapated and anthracnose-resistant accession (95L368), as well as a population of twenty-eight selected recombinant inbred lines (RILs), in a study of the relationship between anthracnose defense and the SGA metabolome. Red fruit tissue extracts of the anthracnose-resistant 95L368 exhibited fungistatic activity against *Colletotrichum* and were found to be enriched in certain SGAs. Correspondingly, gene expression analysis conveyed modulated transcription of GAME genes in 95L368 in both green and red fruit. US28, 95L368, and the descendant RILs displayed variable anthracnose resistance, and metabolomics analysis of this population yielded a distribution of SGA profiles in red fruit. Further analysis of the SGA metabolic profiles identified positive correlations between anthracnose resistance and the α-tomatine and dehydrotomatine derivatives hydroxytomatine, acetoxytomatine, and dehydro-acetoxytomatine. Lastly, Virus-Induced Gene Silencing (VIGS) was employed to evaluate the roles of *GAME31* and *GAME5* in anthracnose resistance, wherein silencing of these GAME genes conferred enhanced levels of anthracnose resistance in a susceptible tomato line. Together, these data provide meaningful insights into the contribution of SGAs to anthracnose defense in tomato, highlighting new targets for the development of crop plants with enhanced biotic disease resistance.

## Materials and Methods

### Plant material and *Colletotrichum* inoculations

The F2 population derived from a cross between the parent lines *Solanum lycopersicum* US28 and 95L368 (formerly denoted 115-4) was generated as described (Stommel, 2001). Twenty-eight recombinant inbred lines (RILs) were selected from a larger RIL population developed via single seed descent from the F2 generation, with each RIL carried to the F6 or F7 generation. Seedlings for US28, 95L368 and the population of 28 RILs were germinated in greenhouse conditions transferred to the field, and arranged into four planting blocks, in Keyport fine loam soil, at USDA-ARS BARC-WEST, Beltsville MD, USA Fruit were harvested at the young, mature green, breaker, and mature red stages, and stored, either whole or diced, at −80 °C for subsequent experiments. For *Colletotrichum* infection assays, mature red fruit from each line (mean n = 22 fruit per block per line) were harvested and transferred to a greenhouse. *Colletotrichum* isolate C9, previously isolated from tomato in Beltsville and tentatively identified as *C. nigrum*, was maintained and prepared as previously described (Stommel and Haynes, 1998). Suspensions were diluted to a final concentration of 5 x 10^6^ spores mL^-1^ for inoculation. Fruit exocarp was wounded via 27-gauge needle, inoculated with a 10 μL droplet of fungal suspension, and incubated in the greenhouse, with lesion diameter (lesion size > 1 mm) recorded at 7 dpi, and where noted, converted to “severity” via normalization to the within-block maximum lesion size.

### *Colletotrichum* growth inhibition assays

Approximately 300 mg red ripe fruit tissue from 95L368 samples stored at −80 °C was ground with liquid nitrogen in a mortar and pestle, left to thaw, then centrifuged at 13,000 rpm for one minute, after which the supernatant was collected. Supernatant was vortexed and centrifuged with equal parts sterile water and 100% MeOH, and samples of whole supernatant, MeOH fraction, water fraction, and MeOH (control) were spread on V8 agar plates. Spores of *Colletotrichum* C9 were harvested from V8 agar plates and suspended in chilled, sterile water, at a final concentration of 5 x 10^3^μL. Aliquots of 10 μL of fungal suspension were added to the PDA plates and incubated at 25 °C for five days for observation of mycelial growth. For the TLC bioassay, 300 mg samples of lyophilized ripe fruit tissue were homogenized in liquid nitrogen, transferred to 10 mL 100% MeOH, and shaken at 250 rpm for 30 minutes. After centrifugation at 5,000 rpm for 15 minutes, 4 mL supernatant was added to 6 mL water, then passed through pre-conditioned (6 mL 100% MeOH followed by 6 mL water) C18 Sep-Pak Cartridges (Waters Corporation, Milford MA, USA) by gravity. After washing with 6 mL 100% MeOH, the samples were eluted with 15 mL 80% MeOH, dried via rotary evaporator, and resuspended in 3 mL 100% MeOH. Aliquots of 10 μL were spotted onto TLC plates and developed at room temperature. After drying the plates in a fume hood for 1 hour, the plates were photographed under UV light, then sprayed with a suspension of *Colletotrichum* C9 (3 x 10^6^ spores mL^-1^) in sterile water and stored in closed petri dishes for 4-5 days before observation of mycelial growth. For the growth inhibition assay incorporating US28, 95L368, and the 4 select RILs, non-treated V8 agar plates were inoculated with *Colletotrichum* as described, and at 7 dpi, 80 μL of MeOH extractions were spotted along the plate perimeters, after which images were generated at 3 and 4 days post-treatment (dpt).

### SGA Metabolomics

For MALDI-TOF MS, fruit tissue methanol extractions separated by C18 chromatography were prepared as previously described. Mass spectra were acquired in positive ion detection mode on a Bruker-Daltonics Microflex spectrometer (Bruker-Daltonics, Billerica MA, USA) For each sample, 2 μL of eluate was mixed with matrix (2,5-dihydrobenzoic acid) in 10 μL acetonitrile and added to 10 μL water, after which 0.5 μL spots of the eluate-matrix mix were added to the stainless-steel target surface and crystallized at room temperature. Each sample was measured in triplicate, with 3,000 laser pulses (λ = 337.1 nm) at 200 ns each, and with the following voltage settings: ion source 1 19.0 kV; ion source 14.0 kV; lens 9.2 kV; and reflector 20.0 kV. Data was processed via the supplied Bruker Daltonics software, and accurate masses were verified via SIS Exact Mass Calculator (https://www.sisweb.com/referenc/tools/exactmass.htm).

For UHPLC-HRMS full scanning, fruit tissue methanol extractions separated by C18 chromatography were prepared as previously described. The UHPLC-HRAM MS system consisted of an LTQ Orbitrap XL MS with an Agilent 1290 UHPLC (Agilent Technologies, Santa Clara CA, USA). Separation was conducted on a Hypersil Gold AQ RP-C18 UHPLC column (Thermo Fisher Scientific, Waltham MA, USA) with an UltraShield pre-column filter (Analytical Scientific Instruments, Richmond CA, USA). Gradient elution (Solution A, 0.1% v/v formic acid in water; Solution B, 0.1% formic acid in acetonitrile) was conducted at a flow rate of 0.3 mL·min^-1^. The linear gradient was as follows: increase from 2% to 10% Solution B (v/v) at 10 minutes; increase to 40% Solution B at 25 minutes; increase to 75% Solution B at 30 minutes; increase to 90% Solution B at 50 minutes; and hold at 90% Solution B to 60 minutes. The re-equilibration time for the initial gradient was 5 minutes. Column compartment temperature was set at 50 °C. Data were acquired under positive ionization mode and negative ionization mode with mass range of *m/z* = 100-2,000 with a resolution of 30,000, separately. The optimized conditions for positive ionization mode were set as follows: sheath gas 80 AU; aux and sweep gas 15 AU; spray voltage 4.5 kV; capillary temperature 325 °C; capillary voltage 40 V; tube lens voltage 150 V; FTMS AGC target 5 x 10^5^; FT-MS/MS AGC target 2 x 10^5^; isolation width 1.5 atomic mass units (amu); and max ion injection time 200 ms. For negative ionization mode, the parameters were set as follows: spray voltage 4 kV; capillary voltage −50 V; and tube lens voltage −120 V. The most intense ion was selected for the data-dependent scan to observe their MS2 to MS4 product ions, respectively, with a normalization collision energy at 35%.

For targeted analysis of SGAs via UHPLC-HRMS, samples of 100 mg lyophilized red fruit tissue were ground in a bead homogenizer with 1/4”-diameter ceramic spheres (MP Biomedicals, LLC, Irvine CA, USA) and vortexed in 5 mL 100% MeOH for 10 seconds. Samples were shaken for 30 minutes at 250 rpm, then centrifuged for 15 minutes at 5,000 rpm, at room temperature. For each sample, approximately 4 mL supernatant was passed through 0.45 μM PTFE syringe filters, then stored at −20 °C. Injection volumes of 1 μL were loaded onto a Q-Exactive mass spectrometer with a Thermo Fisher Vanquish UHPLC system. Separation was conducted on an Agilent Poroshell HPH-C18 column. Gradient elution (Solution A, 0.1% v/v formic acid in water; Solution B, 0.1% v/v formic acid in acetonitrile) was conducted with a flow rate of 0.3 mL ·min^-1^, a sample compartment temperature of 5 °C, and a column temperature of 40 °C. Mass spectra were generated via electrospray ionization with 4 kV spray voltage, an ion transfer tube temperature of 320 °C, and a scan range of m/z 120-1,800 Da. Ion chromatograms were constructed for the protonated molecular ions of each analyte via high resolution ion extraction with a mass width of 0.01 amu, and the SGA contents (μg g dry weight (DW)^-1^) for all analytes were semi-quantitated as α-tomatine equivalent from an external calibration curve of α-tomatine standard (Sigma-Aldrich, St. Louis MO, USA). For statistical analyses, the SGA content values were log10-transformed via the following formula: log10-SGA = log10 [(x μg gDW^1^)+1)].

### Gene expression analysis

Fresh-frozen or lyophilized samples of approximately 100 mg fruit tissue were ground in liquid nitrogen in a mortar and pestle, or via bead homogenizer with 1/4”-diameter ceramic spheres (MP Biomedicals, LLC, Irvine CA, USA). RNA was purified via RNEasy Mini Kit (Qiagen, Hilden DE), per manufacturer’s instructions, with elutions in 30 μL RNase-free water. cDNA synthesis was conducted via ProtoScript II First Strand cDNA Synthesis Kit (New England Biolabs, Ipswich MA, USA), using manufacturer’s instructions for reverse transcription of total RNA. RNA masses and purities were analyzed via a DeNovix DS-11 spectrophotometer (DeNovix Inc., Wilmington DE, USA). cDNA samples were diluted 100-fold for qRT-PCR. Primers for qRT-PCR were designed from target mRNA sequences, spanning introns where possible, and synthesized by Integrated DNA Technologies, Inc. (Coralville IA, USA); see Table S3 for a list of all primer sequences. Primers for GAME-VIGS qRT-PCR validation were designed to yield amplicons without overlap to the VIGS target amplicons (see below). Primer efficiencies, specificities, and optimal annealing temperatures were analyzed via CFX Maestro Software Version 2.2 (Bio-Rad). qRT-PCR was carried out via an iProof SYBR Green Supermix kit (Bio-Rad Laboratories, Hercules CA, USA) per manufacturer’s instructions. Two technical replicates were included for each sample. For each target relative expression (RE) was calculated via the 2^-ddCt^ method, with the lowest-expression sample designated as the calibrator.

### Virus-Induced Gene Silencing (VIGS) cloning and GAME-VIGS Agroinfiltration

RNA and cDNA were isolated from fresh-frozen 95L368 fruit tissue samples as described above. Primers designed to amplify a 406-bp region of *GAME31* coding sequence (CDS), as well as a 478-bp region of *GAME5* CDS, were synthesized by Integrated DNA Technologies and are listed in Table S3. Amplification of target amplicons with the addition of 5’ XbaI and 3’ BamHI restriction enzyme digestion sites was carried out on a Bio-Rad T100 thermocycler (Bio-Rad Laboratories, Hercules CA, USA) via Bio-Rad iProof High Fidelity DNA Polymerase using manufacturer’s instructions. Amplicons were separated via gel electrophoresis on a Bio-Rad PowerPac Basic system, analyzed via an Azure c150 Gel Imaging System (Azure Biosystems, Inc., Dublin CA, USA), and purified via Zymoclean Gel DNA Recovery kit (Zymo Research, Irvine CA, USA) using manufacturer’s instructions. The bipartite VIGS system consisted of the vectors pYL192 (TRV1) and pYL156 (TRV2). Double digestion of pYL156 and target amplicons was conducted via Thermo Fisher FastDigest BamHI and XbaI using manufacturer’s instructions. Ligations utilized New England Biolabs T4 ligase and were conducted at 16 °C overnight, followed by enzyme deactivation at 65 °C for 10 minutes. Aliquots of 2 μL of each reaction were added to 50 μL high efficiency New England Biolabs 5-alpha Competent *E. coli*, placed on ice for 30 minutes, incubated in a 42 °C water bath for 40 seconds, then returned to ice for 2 minutes. After recovery in 1 mL LB for 1 hour, cells were washed in 100 μL LB, plated on LB agar supplemented with 50 μg/mL kanamycin, and incubated at 38 °C overnight. Positive clones were identified via PCR as previously described, using MangoMix Taq DNA polymerase (Meridian Bioscience Inc., Cincinnati OH, USA) and manufacturer’s instructions.

Putatively recombinant plasmid was isolated from overnight cultures using the Zymo Research ZymoPURE Plasmid Miniprep Kit, per manufacturer’s instructions. Positive recombinant vector was verified via Sanger sequencing incorporating the NOS-R primer (Table S3) and performed by Psomagen (Rockville MD, USA). Aliquots of 1 μL of TRV1 and recombinant TRV2 were electroporated into 50 μL tubes of *Agrobacterium tumifasciens* strain GV3101 cells (Intact Genomics, St. Louis MO, USA) using a Bio-Rad Gene Pulser II and Pulse Controller electroporation system set to 2.2 kV. Transformants were grown on LB plates supplemented with 50 μg mL^-1^ kanamycin and 25 μg mL^-1^ spectinomycin and verified via PCR; see Table S3 for primers. For agroinfiltration, 5 mL overnight cultures in LB supplemented with 50 μg mL^-1^kanamycin were transferred to 20 mL cultures and shaken at 225 rpm for 3 hours or until an approximate OD_600_ of 1.0 was reached, then pelleted and resuspended in 20 mL infiltration medium (200 μL 1M MgCl2 + 200 μL 1M MES pH 5.6 + 20 μl 200 mM acetosyringone + 19.58 mL sterile water). After gentle shaking for 3 hours, 1:3 TRV1:TRV2 mixed suspensions were prepared. Micro Tom tomato plants grown in greenhouse conditions and presenting immature green fruit were agroinoculated via the Agrodrench method, in which 5 mL of 1:3 TRV1:TRV2 suspensions were aliquoted to root crowns (Ryu et al., 2004). At 22 dpi, this process was repeated for a second dose of GAME-VIGS Agrodrench. At 41 days following the initial dose, mature red tomatoes were harvested for *Colletotrichum* C9 inoculation and lesion size measurement as previously described.

### Data analysis and software

Statistical significance was determined via one-way Student’s T-test or ANOVA with post-hoc Tukey testing, as noted, using Microsoft Excel or R Studio version 2021.09.0 with R version 4.1.1, with the “multcompView” package, as appropriate. Bar plots were generated in R with the “ggpubr” package. Correlograms were generated, including determination of Pearson correlation coefficients, in R via the “ggally” package. Scatterplots were generated in R via the “ggplot2” package. Heatmaps were generated in R via the “pheatmap” package. Plasmid maps were generated in Geneious Prime version 2022.0.1.

## Results

### Tissue extracts of an anthracnose-resistant, wild tomato accession convey fungicidal activity against *Colletotrichum*

To evaluate the respective levels of anthracnose resistance in US28 and 95L368, ripe fruit were harvested, inoculated with *Colletotrichum*, and scored for lesion diameter at 9 dpi (Fig. S2a). We observed characteristic anthracnose lesions in US28 fruit, however 95L368 fruit were minimally symptomatic and in total, displayed a 33-fold decrease in lesion size when compared to US28 (Fig. 1a-b). We sought to determine whether the enhanced anthracnose resistance exhibited by ripe 95L368 fruit was the result of antifungal, biochemical activity in fruit tissue. Utilizing whole supernatant, water-extracted, and methanol-extracted fruit tissue samples supplemented onto PDA plates, we observed a restriction of mycelial growth on plates supplemented with whole supernatant or methanol-extracted fraction, indicative of fungistatic activity of polar compounds that accumulate in 95L368 fruit (Fig. 1c). To further validate these findings, methanol extractions of US28 and 95L368 ripe fruit were separated via TLC, after which a suspension of *Colletotrichum* C9 (3 x 10^6^ spores mL^-1^) was sprayed onto the TLC plate. At 3-4 dpi, a zone of inhibition around the polar phase of 95L368 fruit tissue extract was observed (Fig. S2b). Together, these findings suggested that one or more polar compounds that accumulate in 95L368 red fruit possess fungistatic activity against *Colletotrichum*, thereby contributing to anthracnose resistance.

**Figure 1.**
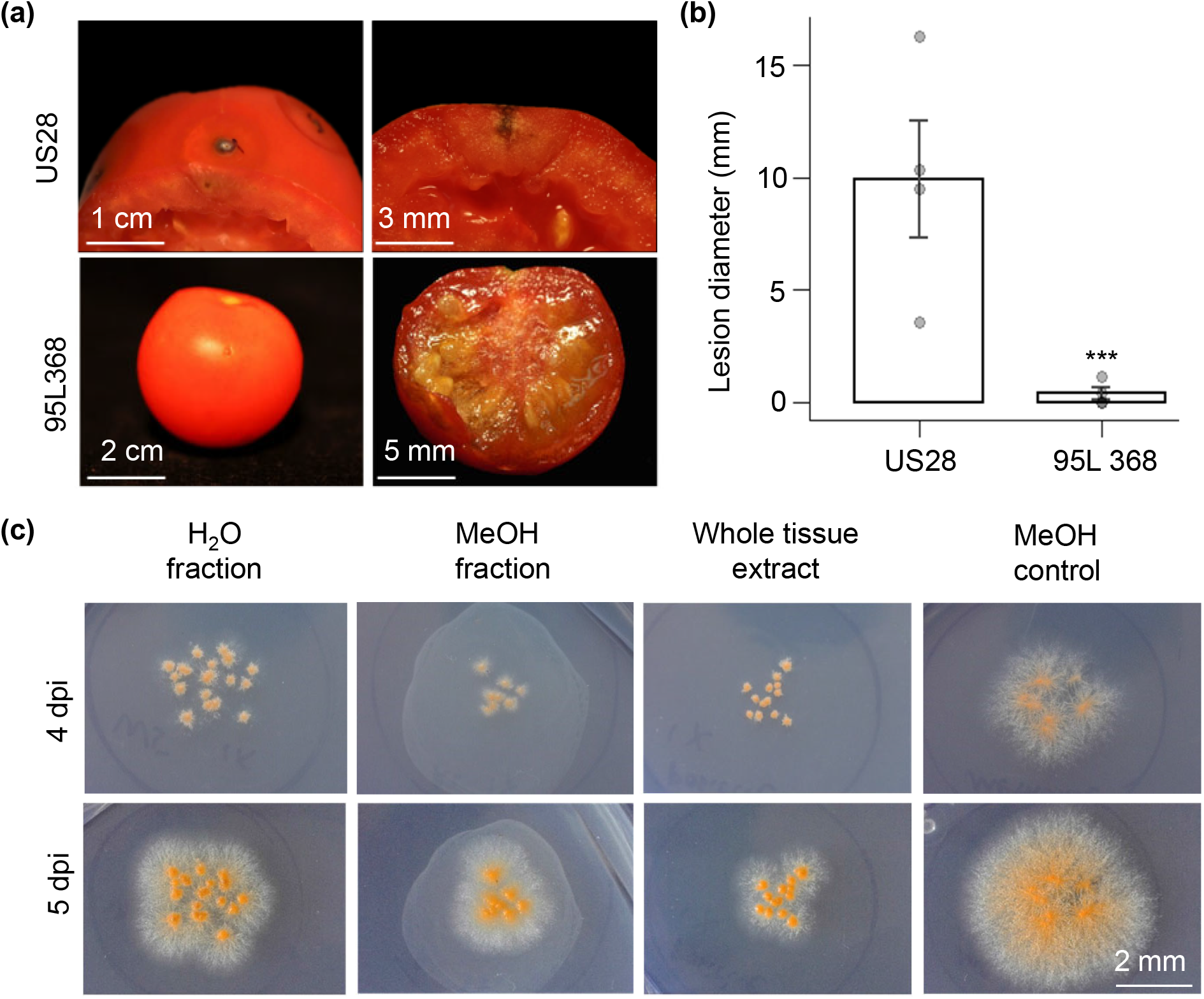
Fruit tissue extracts from an anthracnose-resistant tomato accession exhibit fungistatic activity against *Colleotrichum*. (a) Representative images (9 dpi) of anthracnose-susceptible (US28, top) and resistant (95L368, bottom) tomato fruit following inoculation with *Colletotrichum*. (b) Lesion diameter following *Colletotrichum* infection. Ripe fruit were harvested and inoculated with *Colletotrichum*, with lesion size measured at 9 dpi. Data points represent average lesion size from each of 4 planting blocks per line, with error bars representing SEM. Asterisks represent statistical significance (one-way Student’s T-test). (c) *Colletotrichum* growth inhibition plate assay. H_2_O- or MeOH-fractionated and whole extracts from 95L368 fruit tissue were spotted on PDA plates before inoculation with *Colletotrichum*.

### Ripe fruit tissue of the anthracnose-resistant accession 95L368 is enriched in the production of α-tomatine and several α-tomatine derivatives

To identify compounds that contribute to enhanced anthracnose resistance in 95L368, we conducted a pilot study of phase-separated polar extracts of ripe US28 and 95L368 red fruit tissue. Tissue extracts were separated via TLC as described, and the regions of the TLC plate corresponding to the zone of inhibition of *Colletotrichum* growth (Fig. S2b) were analyzed via HPLC-MS. In a comparison of the total ion chromatograms (TICs) generated for 95L368 and US28 samples we identified elution of several steroidal glycoalkaloids (SGAs) in the retention time range of 20-25 minutes (Fig. 2a; Fig. S3). Additionally, TICs of US28 red fruit displayed a peak at 19.9 minutes that was identified as esculeoside A *(m/z* = 1,270.61). Notably, this peak was absent in TICs of 95L368, which instead displayed strong peaks at 23.1 and 22.7 minutes, indicative of an enrichment of α-tomatine (*m/z* = 1,034.55) and the derivative hydroxytomatine (*m/z* = 1,092.56), respectively.

**Figure 2.**
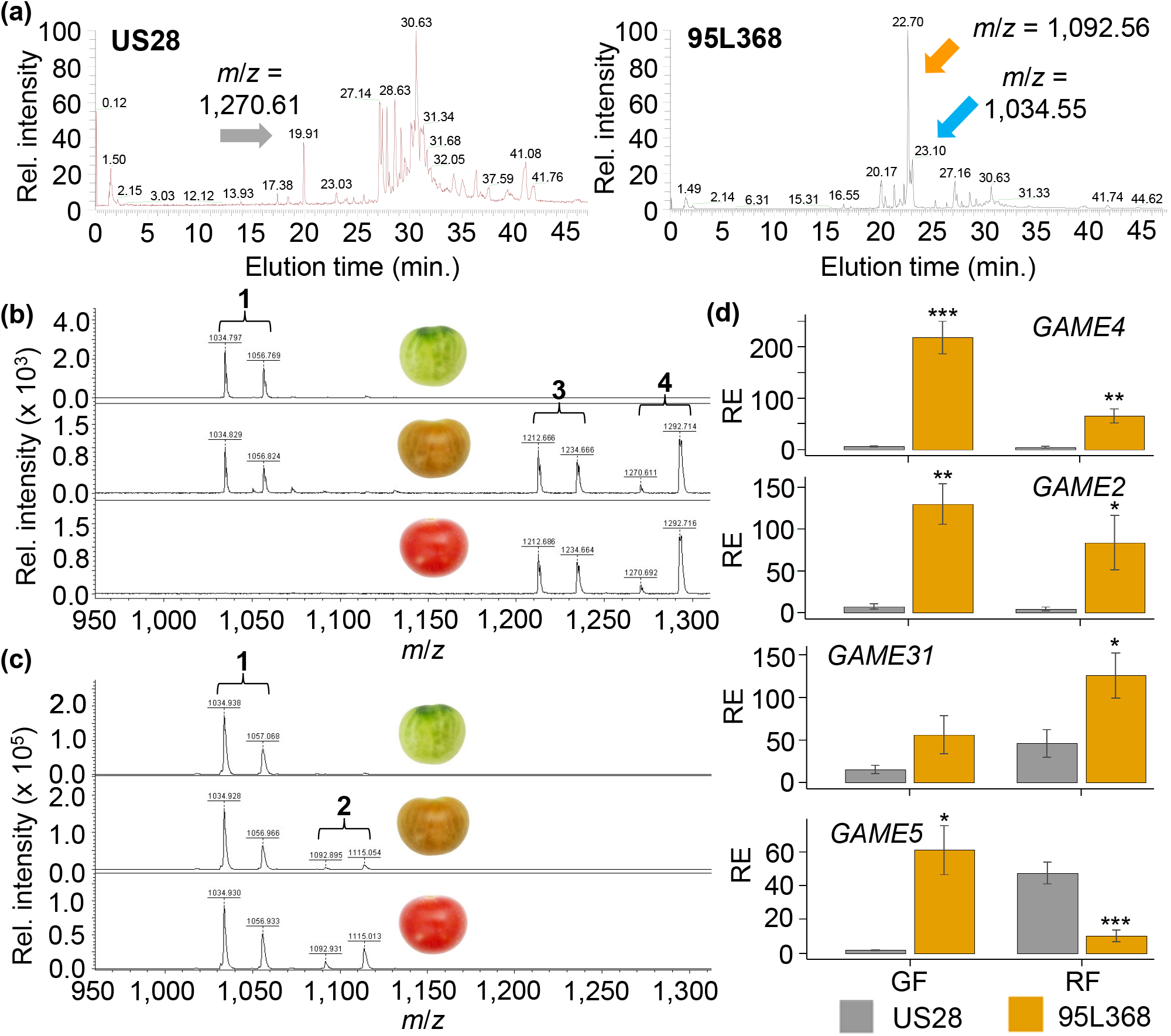
SGA metabolic and *GAME* transcript profiles in green and ripe red fruit of anthracnose-susceptible and –resistant tomato cultivars. (a) Representative TICs from electrospray negative ionization HPLC-MS of ripe red fruit tissue samples of US28 (left) and 95L368 (right). Arrows indicate peaks corresponding to esculeoside A (grey), α-tomatine (orange), and hydroxytomatine (blue). (b) and (c) Representative chromatograms from MALDI-TOF MS analysis of US28 (b) and 95L368 (c) fruit tissue by ripening stage. From top to bottom: green fruit, breaker fruit, and red fruit. From left to right: (1) α–tomatine; (2) acetoxytomatine; (3) unknown compound; and (4) esculeoside A. (d) Expression analysis of GAME gene expression in US28 and 95L368 in green (GF, left) and red (RF, right) fruit. qRT-PCR was conducted with 2 technical replicates of 3 biological replicates per sample, in 3 experiments. RE indicates relative expression, asterisks indicate statistical significance (1-way Student’s T-test), and error bars represent SEM.

We next sought to evaluate the distribution of several key SGAs in green, breaker (intermediate ripening), and ripe red fruit. Methanol extractions of tissue samples from US28 and 95L368 fruit were analyzed via matrix-assisted desorption/ionization time-of-flight (MALDI-TOF) mass spectrometry. In the resulting spectrogram for US28, we observed signals (*m/z* ≈ 1,034 and 1,056), indicative of an accumulation of α-tomatine (C_50_H_83_NO_21_) and its sodium adduct ion in both green and breaker fruit (Fig. 2b). As expected, these α-tomatine signals were also detected in breaker fruit, but not in red ripe fruit, while both breaker and red fruit yielded signals (m/z ≈ 1,270 and 1,292) corresponding to the major and sodium adduct ions of esculeoside A, consistent with the working model for SGA biochemistry in ripening tomato fruit. Interestingly, an unidentified compound yielded suspected major (m/z ≈ 1,213) and sodium adduct (m/z ≈ 1,236) ions, perhaps indicative of deacetoxylated esculeoside A or dehydroxylated esculeoside B (C_56_H_92_NO_27_; MW = 1,211.32), in both breaker and red fruit samples from US28. Analysis of the spectra for green and breaker fruit tissue of 95L368 identified an accumulation of α-tomatine, however, in contrast to US28, α-tomatine was also detected in ripe fruit tissue (Fig. 2c). Additionally, esculeosides were not observed in the spectra for breaker and red fruit samples of 95L368; instead, these samples yielded strong signals (m/z ≈ 1,093 and 1,115) identified as the major and sodium adduct ions for acetoxytomatine, the most proximal derivative of α-tomatine in the SGA metabolic pathway.

### GAME genes are variably expressed in anthracnose-susceptible and -resistant cultivars

Because we observed that fruit of the anthracnose-resistant 95L368 accumulated α-tomatine and hydroxytomatine in both green and red fruit, whereas these compounds decline in ripe fruit of anthracnose-sensitive US28, giving way to the derivative esculeosides, we surveyed the expression of GAME genes, which encode enzymes in the SGA biosynthetic pathway, in both green and red fruit of US28 and 95L368. Overall expression of *GAME4* and *GAME2* declined in the transition from green to red fruit, and in a comparison to US28 green fruit, we observed 95L368 green fruit to exhibit 37.4- and 17.7-fold higher expression of *GAME4* and *GAME2*, respectively (Fig. 2d). Interestingly, red 95L368 fruit were characterized by 2.7-fold higher expression of *GAME31*, as well as a 4.6-fold depletion of *GAME5* transcripts, when compared to red US28 fruit. This expression profile of GAME genes in anthracnose-resistant 95L368 supports our observation of enhanced accumulation of α-tomatine and acetoxytomatine in both green and red fruit, as well as the depletion of esculeosides in red ripe fruit.

### SGA metabolic profiles correlate with anthracnose resistance in RILs developed from a cross between US28 and 95L368

Previous work in our laboratory utilized a tomato population developed from crosses between US28 and multiple different anthracnose-resistant breeding lines, including 95L368, to characterize the heritability of anthracnose resistance (Stommel, 2001; Stommel and Haynes, 1998). Because anthracnose resistance was determined to be a polygenic trait, we evaluated a population of twenty-eight recombinant inbred lines (RILs), developed from a cross between US28 and 95L368, that exhibit variety in anthracnose susceptibility. This population, as well as the parent lines US28 and 95L368, were surveyed for *Colletotrichum* resistance, yielding a distribution of mean lesion diameters ranging from 0.39 to 11.33 mm (Fig. 3; Table S1). Next, we profiled the SGA metabolomes of this population. Methanol extractions separated and analyzed via UHPLC-HRMS yielded mean total SGA levels between 137.21 and 58,929.66 μg g dry weight^-1^, with peaks corresponding to α-tomatine, as well as peaks putatively identified as five SGA derivatives of α-tomatine: multiple isomers of hydroxytomatine *(m/z* = 1,050.55); acetoxytomatine (*m/z* = 1,092.56); acetoxy-dehydrotomatine (*m/z* = 1,090.54); esculeoside A (*m/z* = 1,270.61); and esculeoside B (*m/z* = 1,228.60) (Table S2; Fig. S4). To evaluate the potential contributions of individual SGAs to anthracnose resistance, we performed correlation analyses on anthracnose susceptibility and SGA levels. Due to variance in overall anthracnose disease severity across planting blocks (Fig. S2; Table S1), lesion diameter values in each planting block were transformed to “severity” values by normalizing to the maximum lesion size in that block. Pearson correlation coefficients for severity and the log10-transformed SGA measurements conveyed negative correlations (−0.5 ≤ R^2^ ≤ −0.68) between disease severity and the SGAs α-tomatine, hydroxytomatine, acetoxytomatine, and dehydro-acetoxytomatine, with hydroxytomatine levels yielding the strongest correlation (R^2^ = −0.68) (Fig. 4a-b; Fig. S5). Among the SGAs measured, hydroxytomatine, acetoxytomatine, and dehydro-acetoxytomatine exhibited strong positive correlations (0.73 ≤ R^2^ ≤ 0.97) with each other, as did esculeoside A and esculeoside B (R^2^ = 0.69).

**Figure 3.**
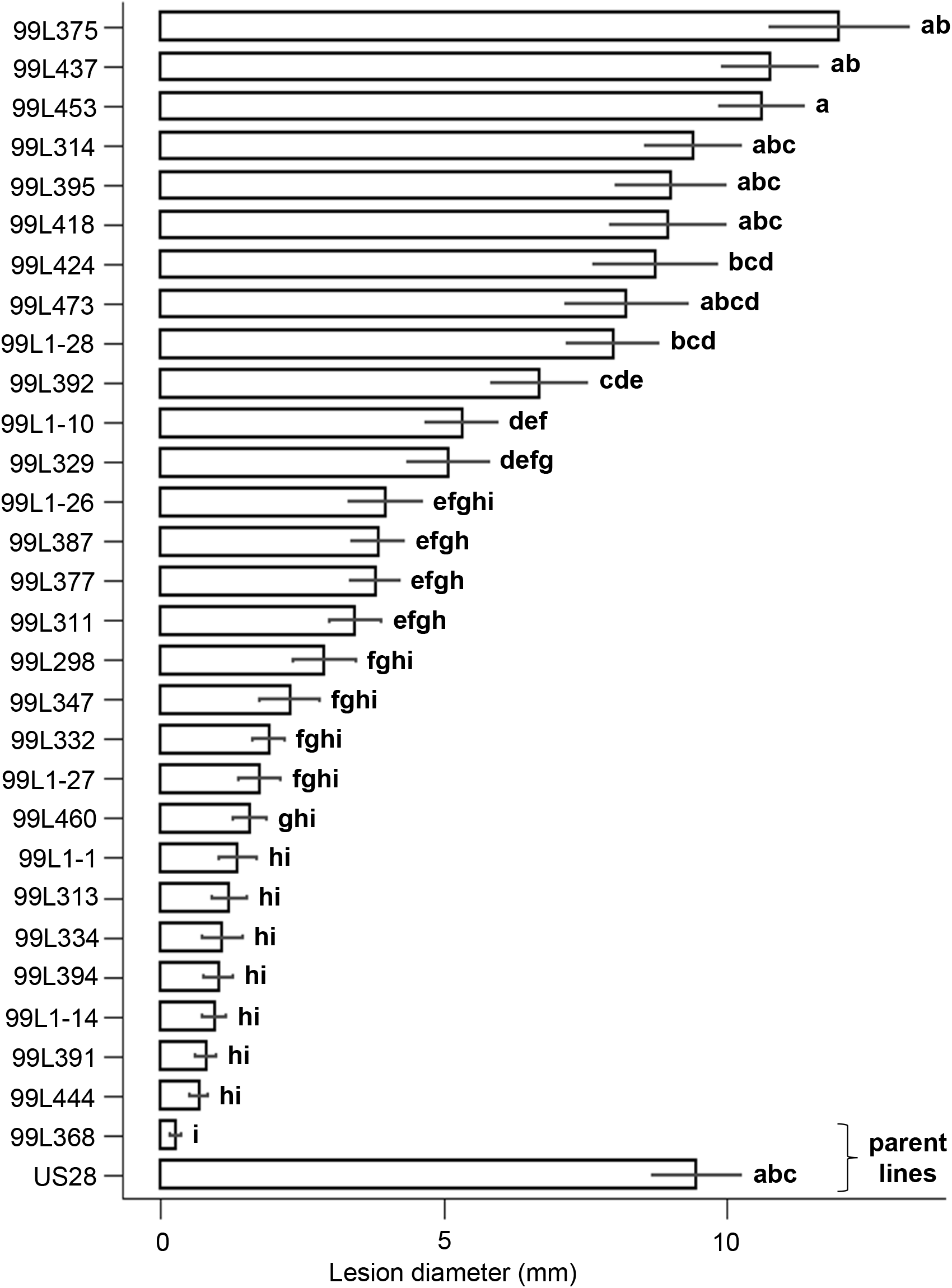
Anthracnose resistance distribution among parent lines and RILs. Ripe red fruit were harvested and inoculated with *Colletotrichum* and incubated in greenhouse conditions, with lesion diameter measured at 9 dpi. Data points represent mean lesion size from each of 4 planting blocks per line, with error bars representing SEM. Statistical significance was determined via ANOVA with post-hoc Tukey testing.

**Figure 4.**
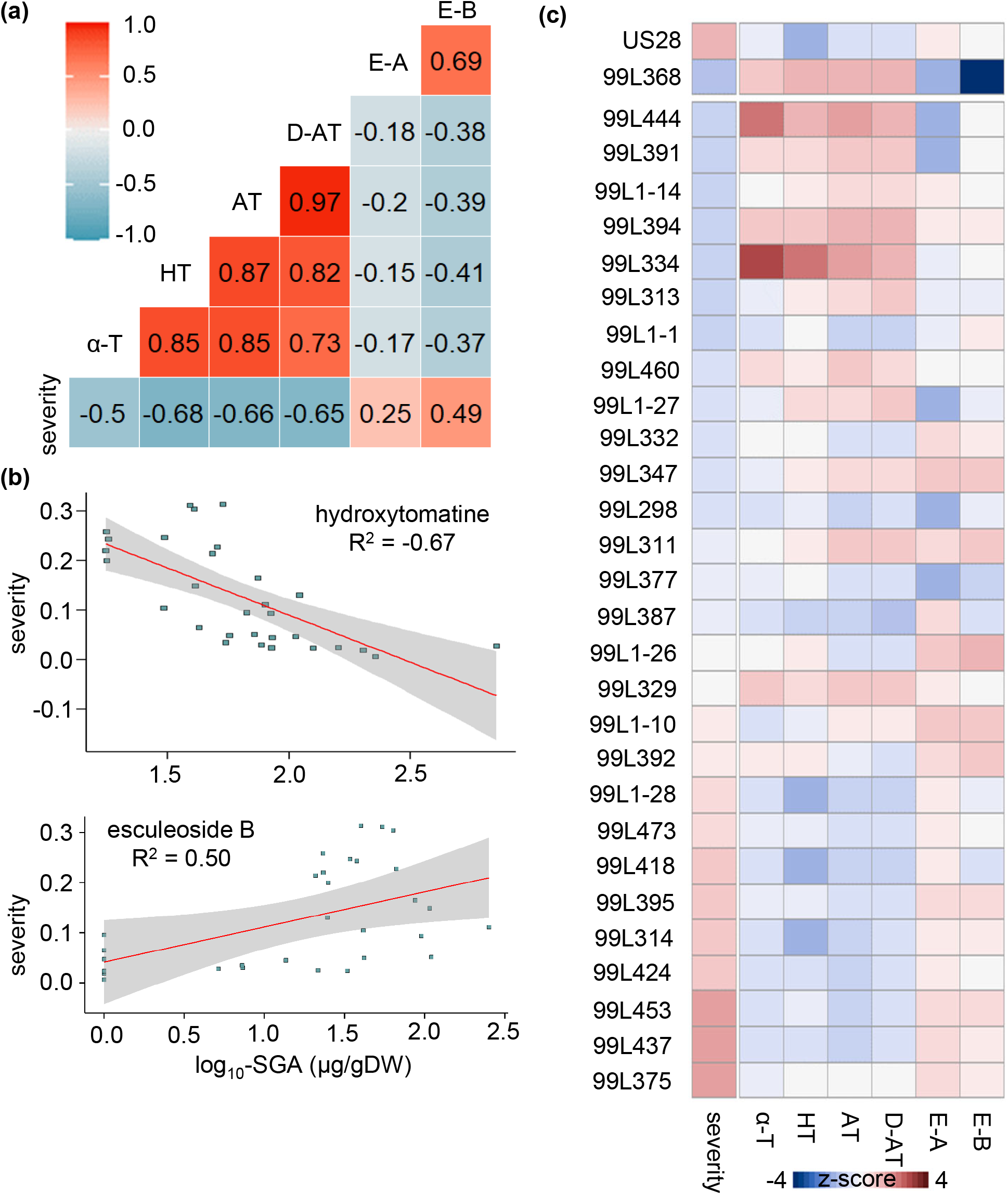
SGAs variably correlate to anthracnose resistance in the parent lines and RIL population. (a) Pearson correlogram of total anthracnose disease severity and SGA levels. (b) Scatterplot of anthracnose severity vs. hydroxytomatine (upper) and esculeoside A (lower) levels with general linear model (GLM) trendline and 95% C.I. Each point corresponds to an individual parent line or RIL. (c) Heat map of z-score-transformed anthracnose severity and SGA levels by parent line and RILs. SGA levels for (a) - (c) are log_10_-transformed. α–T, α -tomatine. HT, hydroxytomatine. AT, acetoxytomatine. D-AT, dehydro-acetoxytomatine. E-A, esculeoside A. E-B, esculeoside B.

To survey the distribution of anthracnose disease severity and accumulation of the individual SGAs at the level of individual RILs and parent lines, we generated a heat map with RILs ordered by increasing anthracnose severity. Interestingly, the six most anthracnose-resistant RILs (99L444, 99L391, 99L1-14, 99L394, 99L334, and 99L313) exhibited a pattern of relative enrichment in hydroxytomatine, acetoxytomatine, and dehydro-acetoxytomatine. Conversely, nine most anthracnose-susceptible RILs (99L375, 99L437, 99L453, 99L424, 99L314, 99L395, 99L418, 99L473, and 99L1-28) were relatively depleted of these analytes and possessed relatively higher levels of the esculeosides (Fig. 4c). Of the SGAs most positively correlated with anthracnose resistance, only hydroxytomatine has been characterized for synthesis by a corresponding GAME gene (*GAME31*). To investigate whether modulated expression of GAME genes is associated with enrichment or depletion of SGAs in RILs, we selected two highly anthracnose-resistant (99L444 and 99L391) and -susceptible (99L453 and 99L375) RILs for GAME gene expression analysis and observed variable enrichment of *GAME2* and *GAME31* transcripts in the selected anthracnose-resistant RILs (Fig. S6a). Additionally, methanol extractions of US28, 95L368, and the four selected RILs were tested in a plate growth inhibition assay using *Colletotrichum* strain C9, wherein methanol extractions from 95L368 and the two anthracnose-resistant RILs displayed inhibitive activity, restricting mycelial growth at the perimeter of the plate, in contrast to the extractions from anthracnose-susceptible lines, which displayed no effect (Fig. S6b).

### Targeted silencing of *GAME31* or *GAME5* confers enhanced anthracnose defense

Following our observations that tomato fruit with enhanced anthracnose resistance are enriched in α-tomatine, hydroxytomatine, acetoxytomatine, and dehydro-acetoxytomatine, we theorized that targeted gene silencing of select GAME genes would improve resistance to *Colletotrichum* in fruit of susceptible wild type tomato. To that end, we employed a bipartite Virus-Induced Gene Silencing (VIGS) system.

Hypothesizing that silencing of *GAME31* would attenuate conversion of α-tomatine to hydroxytomatine, thereby increasing α-tomatine levels, and that silencing of *GAME5* would reduce conversion of SGAs to the esculeosides, we cloned a 406-bp region of *GAME31* coding sequence (CDS) and a 478-bp region of *GAME5* CDS into the pYL156 TRV2 plasmid (Fig. S7). The resulting TRV1-TRV2 vector systems, in addition to a TRV1-TRV2 empty vector system, were introduced to Micro Tom plants with developing fruit via the Agrodrench method (Ryu et al., 2004). Ripe fruit were harvested for *Colletotrichum* infection, and at 7 dpi, we observed 4.3-fold and 3.4-fold reductions in mean lesion diameter in *GAME31-VIGS* and *GAME5-*VIGS transformant fruit, respectively, accompanied by 3.6-fold and 2.4-fold reductions in target gene expression (Figure 5).

**Figure 5.**
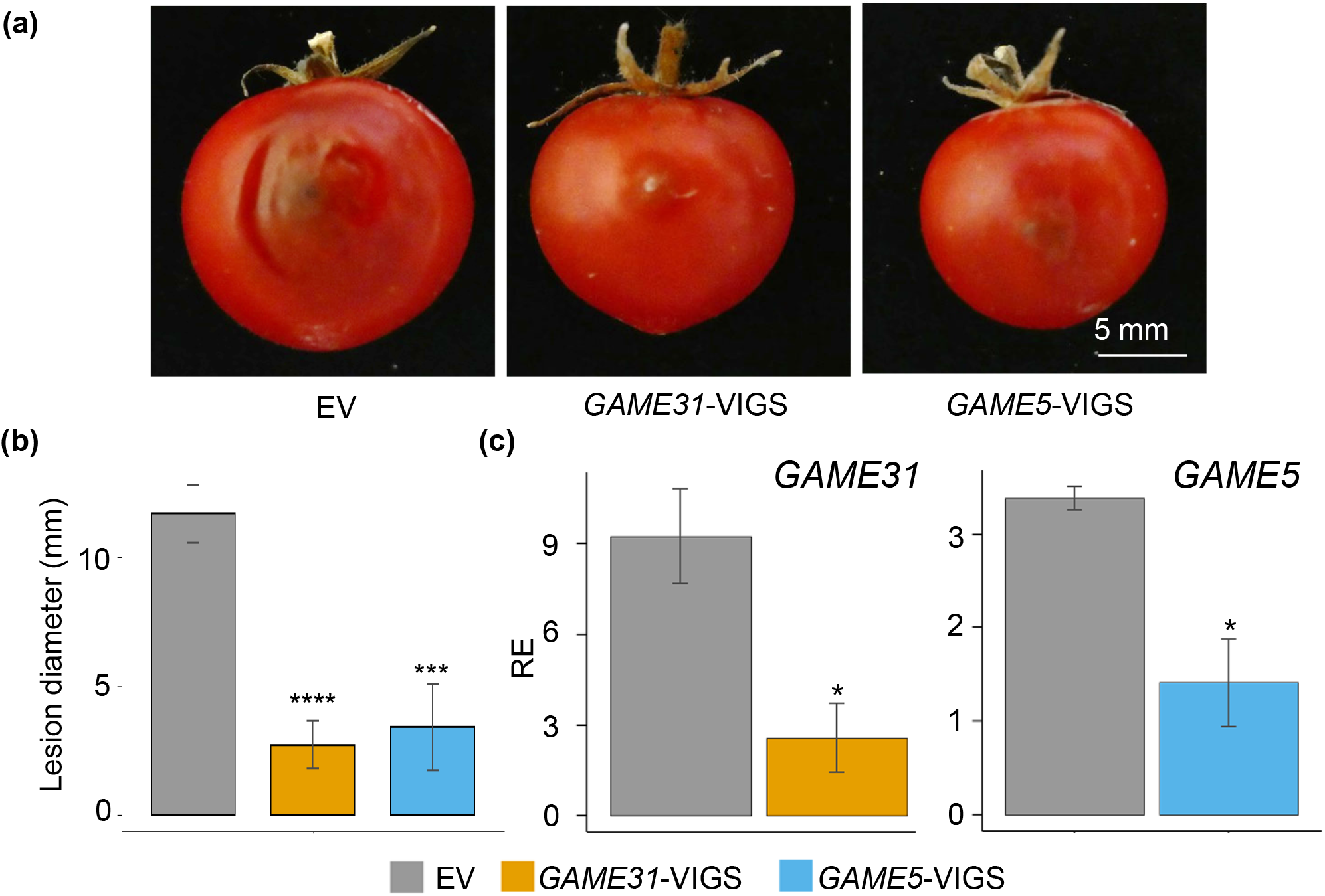
Transient, targeted silencing of *GAME31* and *GAME5* confer enhanced anthracnose resistance in wild type fruit. Bipartite VIGS vectors targeting silencing of *GAME31* or *GAME5* were introduced to Micro Tom plants (n = 2 per treatment) presenting immature green fruit via the Agrodrench method, and mature red fruit were harvested for *Colletotrichum* infection and gene expression via qRT-PCR. (a) Representative images of infected fruit. (b) Lesion diameter of infected fruit (n > 9). (c) Expression of *GAME31* (left) and *GAME5* (right). Samples from infected fruit were collected at 7 dpi and processed for RNA extraction. For each target, qRT-PCR was conducted with 2 technical replicates of 3 biological replicates per treatment. EV, empty vector control. RE indicates relative expression, asterisks indicate statistical significance (* p < 0.05, *** p < 0.001, **** p < 0.0001, 1-way Student’s t-test) and error bars indicate SEM.

## Discussion

Anthracnose, caused by the hemibiotrophic fungal pathogen *Colletotrichum spp*., affects a wide array of crop plants in the field and post-harvest, negatively impacting yield and threatening food security in many growing regions across the globe. To date, there have been limited studies providing insights into mechanisms of anthracnose resistance. Here, our work incorporating an anthracnose-resistant tomato breeding line, as well as a population of 28 RILs derived from a cross between anthracnose-resistant and -susceptible parentage, identifies a role for the secondary metabolites known as steroidal glycoalkaloids (SGAs) in anthracnose defense in tomato. Our data illustrates a fungistatic activity against *Colletotrichum* in red ripe fruit tissue extracts of the anthracnose-resistant breeding line 95L368, and through comparative biochemical analysis of 95L368 and the anthracnose-susceptible cultivar US28, an enrichment in ripe 95L368 fruit of the SGAs α-tomatine and acetoxytomatine accompanying an absence of the end-product esculeosides. Incorporating the population of RILs in a broader analysis of anthracnose resistance and SGA metabolomic profiles, we identified positive correlations between anthracnose resistance and the α-tomatine derivatives hydroxytomatine and acetoxytomatine, as well as dehydro-acetoxytomatine, a dehydrotomatine derivative, respectively. Enhanced expression of GAME genes, which encode enzymes in SGA biosynthesis, was observed in 95L368 and select anthracnose-resistant RILs, and targeted silencing of *GAME31* and *GAME5* conferred enhanced anthracnose resistance in fruit of susceptible wild type plants. Together, our findings introduce a role for several SGAs, previously shown to accumulate under developmental control during the ripening process in tomato, in defense against anthracnose. Our data further suggest that the transition of SGA profiles underpins the transition of *Colletotrichum* infections from a quiescent state to necrotrophy during tomato fruit ripening.

Across our population of anthracnose-resistant and -susceptible parent and RIL lines we observed a distribution of SGA accumulation (Fig. 4; Fig. S5; Table S2), as well as modulation of GAME gene expression, namely enhanced expression of *GAME4, GAME2*, and *GAME31*, coincident with enhanced anthracnose resistance (Fig. 3; Fig. S6). Additionally, targeted silencing of *GAME31* and *GAME5* conferred enhanced levels of anthracnose resistance in wild type fruit (Fig. 5; Fig. S7). In tomato fruit, the transition of *Colletotrichum* infections from a quiescent state to necrotrophy coincides with ripening, and our data suggest that favorable variations in GAME gene expression elicit SGA metabolic profiles that enhance anthracnose defense. The nature of the underlying regulation of GAME gene expression remains to be elucidated. However, the ripening process in tomato fruit is characterized by a variety of changes within the host tissue, such as the activation of production and signaling of the phytohormones abscisic acid, ethylene, and jasmonic acid, and accordingly, phytohormones have been characterized for roles in the regulation of SGA biosynthesis in tomato (Alkan et al., 2015; Cárdenas et al., 2015; Pesaresi et al., 2014; Quinet et al., 2019). Research incorporating tomato mutant lines with constitutive production of jasmonic acid, as well as exogenous treatment with methyl jasmonate (MeJA), have yielded enhanced levels of α-tomatine and expression of GAME genes in root and leaf tissue; conversely, leaf tissue from mutant lines with attenuated jasmonic acid production displayed lower levels of α-tomatine and several derivatives, as well as impaired defense against the fungal pathogen *Fusarium oxysporum* (Montero-Vargas et al., 2018; Thagun et al., 2016). Transcription factors known as jasmonate-responsive transcription factors of the ETHYLENE RESPONSE FACTOR (ERF) family (JREs) interact with JRE-binding elements in gene promoters and have been investigated for their role in the regulation of GAME gene expression. *GAME9*, also known as JRE4, is jasmonic acid-induced and expressed at high levels in various plant tissues, and interacts with the JRE-binding element in the 5’ flanking regions of *GAME4* (Thagun et al., 2016; Yu et al., 2020). *JRE4* has also been shown to positively regulate the expression of multiple different GAME genes, and transgenic suppression of *JRE4* in tomato fruit conferred reductions in α-tomatine content in green fruit (Nakayasu et al., 2018; Thagun et al., 2016). Modulation of GAME gene expression, which impacts SGA production and anthracnose defense, may result from variable levels of jasmonic acid signaling and JRE transcription factor activity at key GAME genes. Therefore, to better understand the genetic mechanisms underlying anthracnose defense, future work should seek to assess endogenous jasmonic acid levels, jasmonic acid signaling activity, and GAME gene promoter identity and activity in anthracnose-resistant and -susceptible lines.

Multiple activities explaining the fungistatic activity of SGAs against pathogens have been proposed. Fungistatic activity of tomatidine against the human fungal pathogen *Candida albicans* was shown to occur through the ergosterol pathway, supporting a role for SGAs in membrane disruption of fungal pathogens (Dorsaz et al., 2017). *C. albicans* treated with tomatidine exhibited dose-dependent reductions of ergosterol, as well as inhibition of the sterol methyltransferase Erg6, and mutations in Erg6 were identified from a forward genetics approach utilizing tomatidine-resistant strains. Multiple studies have demonstrated the efficacy of α-tomatine in membrane disruption using liposomes, with reports of specificity of α-tomatine towards 3β-hydroxyl sterols such as ergosterol which, in free form, are present in lower amounts in tomato (Roddick and Drysdale, 1984; Steel and Drysdale, 1988; You and Kan, 2021). Alternatively, the induction of cell death in *Fusarium oxysporum* by α-tomatine was shown to occur via the generation of intracellular reactive oxygen species (ROS), likely through modulation of tyrosine kinase, G protein, and ATP synthase activities (Ito et al., 2007). The tetraose moiety of α-tomatine is important to its biological activity: Tomatinases, produced by various tomato pathogens, are capable of hydrolyzing one or more carbohydrate residues from this moiety, rendering α-tomatine less toxic and thereby contributing to virulence (Ökmen et al., 2013; Pareja-Jaime et al., 2008; Sandrock and VanEtten, 1998). Targeted manipulation of the SGA metabolic pathway and its metabolites may afford opportunities for durable host resistance against pathogen detoxification mechanisms.

We employed VIGS to silence expression of *GAME31* and *GAME5*, respectively targeted to reduce the conversion of α-tomatine to hydroxytomatine and the production of the esculeosides, with both strategies conferring enhanced anthracnose resistance (Fig. 5; Fig. S7). Our results demonstrated fungistatic activities of red tomato fruit tissue extracts possessing high levels of α-tomatine and the derivatives hydroxytomatine, acetoxytomatine, and dehydro-acetoxytomatine, and positive correlations between the accumulation of these compounds and anthracnose resistance in red fruit (Fig. 1–2; Fig. S2-S4; Fig. 4; Table S2). One or more of these derivatives may possess antifungal activity similar to that of α-tomatine, however, no studies have individually examined these α-tomatine derivatives for their antifungal activities. Therefore, future studies might seek to isolate these compounds for structural evaluation of their fungistatic activities against *Colletotrichum* and other fungal pathogens of tomato and other crop species. Furthermore, the targeted manipulation of GAME gene expression presented here can be expanded to develop tomato breeding lines with stable, targeted enhancement or suppression of GAME gene expression, with the purpose of generating high levels of SGAs conferring enhanced resistance to *Colletotrichum* and other tomato pathogens. Together, our data demonstrates a role for SGAs, including α-tomatine and certain of its derivatives, as well as GAME genes including *GAME31* and *GAME5*, in anthracnose defense in tomato fruit, findings that can be utilized to further optimize management of tomato pathogens that impact yield.

## Abbreviations

GAME: glycoalkaloid metabolic enzyme
MALDI-TOF: matrix-assisted laser desorption/ionization time-of-flight
MS: mass spectrometry
SGA: steroidal glycoalkalkoid
UHPLC: ultra-high performance liquid chromatography

## Supplementary Data

Fig. S1. Simplified schematic of the steroidal glycoalkaloid (SGA) metabolic pathway during tomato fruit ripening.

Fig. S2. *Colleotrichum* resistance in the tomato accession 95L368.

Fig. S3. 95L368 accumulates enhanced levels of SGAs.

Fig. S4. Representative extracted ion chromatograms (EICs) for detected SGAs.

Fig. S5. Mean detected SGAs vs. mean anthracnose disease severity.

Fig. S6. Anthracnose-resistant and –susceptible RILs exhibit variable GAME gene expression and *Colletotrichum* growth inhibition.

Fig. S7. Transient silencing of *GAME* genes via VIGS.

Table S1. Anthracnose resistance distribution among parent lines and RILs.

Table S2. SGA measurements (ug gDW^-1^) among parent lines and RILs.

Table S3. PCR primers utilized in this study.

## Acknowledgements

This research was supported by the U.S. Department of Agriculture, Agricultural Research Service. Mention of trade names or commercial products in this publication is solely for the purpose of providing specific information and does not imply recommendation or endorsement by the U.S. Department of Agriculture. USDA is an equal opportunity provider and employer.

## Author Contributions

Fabian, M.L. and Zhang, C. contributed equally to this study.

