## Supplemental Tables 1, 2, 3 for "Steroidal glycoalkaloids contribute to anthracnose resistance in *Solanum lycopersicum*"

**Table S1.** Anthracnose resistance distribution among parent lines and RILs. Ripe red fruit (mean n=22 fruit/line/block) were harvested from each planting block and inoculated with Colletotrichum, with lesion size measured at 9 dpi.

| | block 1 lesion diameter (mm) | | | block 2 lesion diameter (mm) | | | block 3 lesion diameter (mm) | | | block 4 lesion diameter (mm) | | |
|---|---|---|---|---|---|---|---|---|---|---|---|---|
| **line** | **mean** | **sd** | **combined** | **mean** | **sd** | **combined** | **mean** | **sd** | **combined** | **mean** | **sd** | **combined** |
| US28 | 16.29 | 7.85 | 16.29 ± 7.85 | 10.33 | 5.35 | 10.33 ± 5.35 | 9.50 | 4.20 | 9.50 ± 4.20 | 3.57 | 3.06 | 3.57 ± 3.06 |
| 95L 368 | 1.15 | 1.46 | 1.15 ± 1.46 | 0.40 | 1.10 | 0.40 ± 1.10 | 0.00 | 0.00 | 0.00 ± 0.00 | 0.00 | 0.00 | 0.00 ± 0.00 |
| 99L 444 | 1.44 | 1.86 | 1.44 ± 1.86 | 0.15 | 0.65 | 0.15 ± 0.65 | 0.73 | 1.35 | 0.73 ± 1.35 | 0.47 | 1.33 | 0.47 ± 1.33 |
| 99L 391 | 0.60 | 1.46 | 0.60 ± 1.46 | 1.40 | 2.09 | 1.40 ± 2.09 | 0.53 | 1.41 | 0.53 ± 1.41 | 1.00 | 1.58 | 1.00 ± 1.58 |
| 99L 394 | 2.06 | 1.98 | 2.06 ± 1.98 | 1.45 | 2.43 | 1.45 ± 2.43 | 0.52 | 1.33 | 0.52 ± 1.33 | 0.00 | 0.00 | 0.00 ± 0.00 |
| 99L 1-14 | 2.45 | 3.93 | 2.45 ± 3.93 | 0.86 | 1.72 | 0.86 ± 1.72 | 0.13 | 0.60 | 0.13 ± 0.60 | 0.79 | 1.94 | 0.79 ± 1.94 |
| 99L 334 | 1.84 | 4.15 | 1.84 ± 4.15 | 0.60 | 1.20 | 0.60 ± 1.20 | 0.78 | 1.81 | 0.78 ± 1.81 | 0.62 | 1.59 | 0.62 ± 1.59 |
| 99L 313 | 2.04 | 2.65 | 2.04 ± 2.65 | 2.04 | 4.70 | 2.04 ± 4.70 | 0.46 | 1.31 | 0.46 ± 1.31 | 0.26 | 0.85 | 0.26 ± 0.85 |
| 99L 1-1 | 2.96 | 5.37 | 2.96 ± 5.37 | 0.63 | 2.06 | 0.63 ± 2.06 | 0.70 | 2.12 | 0.70 ± 2.12 | 1.45 | 2.04 | 1.45 ± 2.04 |
| 99L 460 | 2.20 | 2.15 | 2.20 ± 2.15 | 0.88 | 2.34 | 0.88 ± 2.34 | 2.05 | 3.32 | 2.05 ± 3.32 | 0.69 | 1.83 | 0.69 ± 1.83 |
| 99L 1-27 | 2.58 | 3.32 | 2.58 ± 3.32 | 1.14 | 1.88 | 1.14 ± 1.88 | 1.33 | 2.36 | 1.33 ± 2.36 | 0.00 | 0.00 | 0.00 ± 0.00 |
| 99L 347 | 2.52 | 2.78 | 2.52 ± 2.78 | 1.33 | 2.19 | 1.33 ± 2.19 | 0.80 | 1.67 | 0.80 ± 1.67 | 2.85 | 3.22 | 2.85 ± 3.22 |
| 99L 332 | 7.89 | 6.32 | 7.89 ± 6.32 | 1.31 | 3.01 | 1.31 ± 3.01 | 0.13 | 0.60 | 0.13 ± 0.60 | 0.55 | 1.95 | 0.55 ± 1.95 |
| 99L 298 | 7.20 | 8.37 | 7.20 ± 8.37 | 1.19 | 2.37 | 1.19 ± 2.37 | 0.93 | 2.16 | 0.93 ± 2.16 | 0.55 | 1.36 | 0.55 ± 1.36 |
| 99L 1-26 | 2.38 | 2.59 | 2.38 ± 2.59 | 1.25 | 2.41 | 1.25 ± 2.41 | 4.97 | 5.59 | 4.97 ± 5.59 | 4.38 | 4.02 | 4.38 ± 4.02 |
| 99L 311 | 7.03 | 4.82 | 7.03 ± 4.82 | 1.30 | 1.78 | 1.30 ± 1.78 | 1.87 | 3.39 | 1.87 ± 3.39 | 4.95 | 4.19 | 4.95 ± 4.19 |
| 99L 387 | 4.50 | 5.48 | 4.50 ± 5.48 | 2.36 | 3.40 | 2.36 ± 3.40 | 2.59 | 3.51 | 2.59 ± 3.51 | 7.05 | 3.94 | 7.05 ± 3.94 |
| 99L 377 | 9.17 | 9.61 | 9.17 ± 9.61 | 2.86 | 4.45 | 2.86 ± 4.45 | 1.32 | 2.77 | 1.32 ± 2.77 | 2.90 | 3.81 | 2.90 ± 3.81 |
| 99L 329 | 7.59 | 7.18 | 7.59 ± 7.18 | 6.71 | 8.29 | 6.71 ± 8.29 | 2.80 | 2.87 | 2.80 ± 2.87 | 3.33 | 4.54 | 3.33 ± 4.54 |
| 99L 1-10 | 5.47 | 6.31 | 5.47 ± 6.31 | 8.00 | 6.73 | 8.00 ± 6.73 | 4.95 | 4.13 | 4.95 ± 4.13 | 2.63 | 2.99 | 2.63 ± 2.99 |
| 99L 392 | 12.50 | 10.28 | 12.50 ± 10.28 | 3.65 | 6.26 | 3.65 ± 6.26 | 2.37 | 4.55 | 2.37 ± 4.55 | 7.45 | 3.77 | 7.45 ± 3.77 |
| 99L 424 | 12.00 | 10.43 | 12.00 ± 10.43 | 3.00 | 3.98 | 3.00 ± 3.98 | 6.93 | 4.63 | 6.93 ± 4.63 | 10.95 | 7.76 | 10.95 ± 7.76 |
| 99L 1-28 | 16.42 | 13.67 | 16.42 ± 13.67 | 4.25 | 4.04 | 4.25 ± 4.04 | 3.48 | 4.61 | 3.48 ± 4.61 | 6.68 | 5.50 | 6.68 ± 5.50 |
| 99L 473 | 15.53 | 11.75 | 15.53 ± 11.75 | 7.22 | 6.02 | 7.22 ± 6.02 | 5.17 | 5.46 | 5.17 ± 5.46 | 5.27 | 5.35 | 5.27 ± 5.35 |
| 99L 395 | 6.85 | 7.87 | 6.85 ± 7.87 | 18.90 | 11.14 | 18.90 ± 11.14 | 5.78 | 6.79 | 5.78 ± 6.79 | 5.32 | 5.16 | 5.32 ± 5.16 |
| 99L 418 | 18.36 | 9.13 | 18.36 ± 9.13 | 5.62 | 4.67 | 5.62 ± 4.67 | 6.20 | 6.83 | 6.20 ± 6.83 | 3.19 | 2.51 | 3.19 ± 2.51 |
| 99L 314 | 14.04 | 8.38 | 14.04 ± 8.38 | 4.52 | 5.05 | 4.52 ± 5.05 | 7.25 | 6.42 | 7.25 ± 6.42 | 11.00 | 7.03 | 11.00 ± 7.03 |
| 99L 437 | 10.71 | 10.25 | 10.71 ± 10.25 | 8.72 | 5.94 | 8.72 ± 5.94 | 9.56 | 5.62 | 9.56 ± 5.62 | 15.18 | 4.06 | 15.18 ± 4.06 |
| 99L 375 | 11.52 | 8.50 | 11.52 ± 8.50 | 4.10 | 6.85 | 4.10 ± 6.85 | 9.87 | 6.14 | 9.87 ± 6.14 | 16.83 | 6.71 | 16.83 ± 6.71 |
| 99L 453 | 23.42 | 13.22 | 23.42 ± 13.22 | 7.27 | 6.54 | 7.27 ± 6.54 | 6.37 | 6.06 | 6.37 ± 6.06 | 8.24 | 4.34 | 8.24 ± 4.34 |
| **Grand Total** | **7.97** | **9.64** | **7.97 ± 9.64** | **3.74** | **6.05** | **3.74 ± 6.05** | **3.36** | **5.10** | **3.36 ± 5.10** | **4.42** | **5.89** | **4.42 ± 5.89** |

**Table S2.** SGA measurements (ug gDW$^{-1}$) among parent lines and RILs. Ripe red fruit (n = 4) were lyophilized and methanol extractions were analyzed via UHPLC-HRMS.

| | α-tomatine | | hydroxytomatines | | acetoxytomatine | | acetoxy-dehydrotomatine | | esculeoside A | | esculeoside B | | total | |
|---|---|---|---|---|---|---|---|---|---|---|---|---|---|---|
| **line** | **mean** | **sd** | **mean** | **sd** | **mean** | **sd** | **mean** | **sd** | **mean** | **sd** | **mean** | **sd** | **mean** | **sd** |
| US28 | 75.42 | 20.63 | 16.65 | 0.16 | 22.76 | 8.79 | 18.31 | 1.83 | 22.06 | 1.16 | 22.24 | 0.91 | 177.44 | 33.26 |
| 95L368 | 241.14 | 144.71 | 226.42 | 50.28 | 6,165.15 | 4,825.69 | 7,361.08 | 3,437.29 | 0.00 | 0.00 | 0.00 | 0.00 | 13,993.79 | 8,359.54 |
| 99L444 | 873.36 | 1,204.34 | 201.15 | 113.63 | 29,766.65 | 43,034.74 | 7,293.42 | 6,559.94 | 39.82 | 40.06 | 20.61 | 35.69 | 38,195.01 | 47,808.40 |
| 99L391 | 167.53 | 81.71 | 124.72 | 34.92 | 2,611.73 | 2,050.38 | 2,783.29 | 2,235.81 | 18.68 | 32.35 | 0.00 | 0.00 | 5,705.95 | 4,335.32 |
| 99L394 | 236.54 | 141.55 | 158.86 | 78.59 | 4,533.81 | 3,774.31 | 3,176.26 | 2,284.56 | 23.53 | 27.86 | 0.00 | 0.00 | 8,129.00 | 6,231.43 |
| 99L1-14 | 99.65 | 54.17 | 83.75 | 18.45 | 680.75 | 672.82 | 686.83 | 378.24 | 22.55 | 13.71 | 0.00 | 0.00 | 1,573.54 | 1,108.05 |
| 99L334 | 1,503.41 | 1,591.33 | 717.06 | 702.16 | 48,330.38 | 54,398.81 | 8,345.63 | 3,662.87 | 23.34 | 40.43 | 4.16 | 7.21 | 58,929.66 | 59,344.04 |
| 99L313 | 88.28 | 24.76 | 75.99 | 55.52 | 671.77 | 654.49 | 1,161.00 | 1,143.39 | 25.81 | 32.35 | 31.88 | 41.88 | 2,054.74 | 1,813.86 |
| 99L1-1 | 71.20 | 5.97 | 53.87 | 36.45 | 20.35 | 2.21 | 13.57 | 7.83 | 88.94 | 73.93 | 108.91 | 74.70 | 370.08 | 193.72 |
| 99L460 | 159.65 | 128.24 | 84.14 | 44.42 | 1,405.24 | 1,617.96 | 1,003.41 | 733.86 | 17.65 | 19.23 | 6.33 | 10.96 | 2,676.40 | 2,453.45 |
| 99L1-27 | 82.65 | 12.36 | 105.36 | 38.26 | 670.46 | 257.79 | 1,187.60 | 449.93 | 19.79 | 20.14 | 0.00 | 0.00 | 2,065.85 | 713.80 |
| 99L347 | 81.64 | 22.64 | 70.93 | 21.63 | 453.69 | 418.14 | 529.01 | 435.99 | 25.84 | 16.19 | 12.61 | 21.84 | 1,173.72 | 880.24 |
| 99L332 | 95.65 | 45.19 | 55.86 | 36.88 | 29.31 | 15.78 | 18.67 | 1.32 | 87.53 | 37.92 | 94.39 | 33.97 | 390.17 | 173.54 |
| 99L298 | 64.17 | 1.38 | 41.54 | 14.07 | 17.30 | 0.40 | 16.72 | 0.15 | 37.93 | 9.20 | 40.89 | 9.77 | 223.02 | 23.74 |
| 99L1-26 | 99.53 | 48.10 | 78.73 | 19.65 | 36.84 | 26.24 | 21.64 | 4.94 | 135.56 | 69.04 | 250.98 | 194.57 | 629.23 | 336.87 |
| 99L311 | 106.16 | 63.39 | 83.13 | 41.87 | 1,284.56 | 1,535.17 | 2,000.43 | 1,895.66 | 35.49 | 27.38 | 6.26 | 10.84 | 3,516.03 | 3,531.19 |
| 99L387 | 66.53 | 6.70 | 29.42 | 14.13 | 18.05 | 2.32 | 4.40 | 7.62 | 13.07 | 22.64 | 40.36 | 13.70 | 171.83 | 65.80 |
| 99L377 | 91.31 | 29.23 | 65.84 | 32.37 | 31.48 | 15.92 | 20.60 | 4.20 | 95.22 | 66.96 | 106.28 | 73.09 | 419.70 | 199.37 |
| 99L329 | 266.45 | 245.25 | 109.54 | 37.42 | 2,803.43 | 2,500.51 | 1,660.28 | 914.88 | 9.23 | 15.99 | 0.00 | 0.00 | 4,848.93 | 3,063.11 |
| 99L1-10 | 71.06 | 9.06 | 40.04 | 31.95 | 254.21 | 270.72 | 353.71 | 399.84 | 28.36 | 22.93 | 23.75 | 28.40 | 779.67 | 665.31 |
| 99L392 | 154.82 | 143.21 | 73.25 | 52.89 | 59.00 | 66.55 | 20.22 | 4.34 | 82.66 | 60.91 | 86.34 | 58.84 | 489.78 | 384.28 |
| 99L424 | 62.53 | 0.29 | 16.54 | 0.19 | 16.95 | 0.56 | 8.61 | 8.61 | 10.28 | 10.38 | 22.30 | 3.53 | 137.21 | 18.75 |
| 99L1-28 | 65.31 | 4.26 | 16.70 | 0.32 | 18.01 | 1.56 | 12.85 | 7.43 | 18.79 | 11.50 | 24.03 | 4.22 | 159.78 | 32.26 |
| 99L473 | 77.37 | 11.82 | 47.32 | 25.34 | 24.38 | 8.29 | 19.39 | 3.81 | 61.29 | 26.20 | 65.67 | 30.98 | 303.78 | 95.63 |
| 99L395 | 63.59 | 0.89 | 17.02 | 0.51 | 18.07 | 0.60 | 17.67 | 0.47 | 26.46 | 3.69 | 19.85 | 12.06 | 162.65 | 12.11 |
| 99L418 | 77.66 | 14.66 | 49.63 | 28.65 | 25.22 | 6.34 | 16.14 | 9.50 | 54.47 | 40.26 | 62.56 | 31.15 | 294.25 | 132.75 |
| 99L314 | 62.86 | 0.71 | 29.67 | 14.06 | 17.74 | 0.81 | 17.57 | 0.78 | 31.94 | 7.67 | 33.20 | 9.23 | 205.76 | 37.30 |
| 99L437 | 70.15 | 7.04 | 38.03 | 13.58 | 20.52 | 2.12 | 18.33 | 1.72 | 34.90 | 3.66 | 36.71 | 4.63 | 227.00 | 8.28 |
| 99L375 | 81.70 | 23.62 | 52.39 | 32.29 | 143.50 | 206.80 | 109.63 | 152.31 | 36.75 | 24.33 | 39.00 | 24.97 | 467.28 | 369.76 |
| 99L453 | 64.00 | 1.05 | 39.60 | 21.28 | 17.54 | 0.61 | 17.00 | 0.59 | 49.98 | 32.08 | 53.48 | 35.11 | 254.36 | 92.12 |
| **Grand Total** | **177.38** | **469.84** | **93.44** | **184.06** | **3,338.96** | **16,156.92** | **1,263.78** | **2,850.99** | **39.26** | **45.08** | **40.43** | **67.93** | **4,957.52** | **18,722.60** |

**Table S3.** PCR primers utilized in this study. Restriction enzyme recognition sites are shown in red.

| primer | sequence (5'-3') | purpose | NCBI target reference | notes | source |
|---|---|---|---|---|---|
| GAME4-F | CAAAGCTTCAATGTGTGGTGATCC` | qPCR amplication of *GAME4* | NM_001365974.1 | | this study |
| GAME4-R | TGCCCTATACATTCCTCCTTCTCC | qPCR amplication of *GAME4* | NM_001365974.1 | | this study |
| GAME2-F | ACGGAAGAAGGTGGTTCCTCTC | qPCR amplication of *GAME2* | NM_001246931.1 | | this study |
| GAME2-R | AGAGCACAACGCTTGATCTCGTG | qPCR amplication of *GAME2* | NM_001246931.1 | | this study |
| GAME31-F | GCCGGTGATTCCTTCAAAGC | qPCR amplication of predicted *GAME31* | XM_004233493.3 | | this study |
| GAME31-R | GCCTGGATGAAGTGACCTGG | qPCR amplication of predicted *GAME31* | XM_004233493.3 | | this study |
| GAME5-F | AGGATCATCTGTCGCGACT | qPCR amplication of *GAME5* | NM_001361347.1 | | this study |
| GAME5-R | TGTGTACCGGGCCTATTGG | qPCR amplication of *GAME5* | NM_001361347.1 | | this study |
| TUB-F | TGGCTACCATCAAGACTAAGCGCA | qPCR amplification of tubulin alpha chain gene | NM_001347592.1 | qPCR reference gene | this study |
| TUB-R | AGACCTCAGCAACACTGGTTGAGT | qPCR amplification of tubulin alpha chain gene | NM_001347592.1 | qPCR reference gene | this study |
| EF-1-F | GATTGGTGGTATTGGAACTGTC | qPCR amplification of predicted EF-1α gene | XM_004240531.4 | qPCR reference gene | Roternberg et al., 2006 |
| EF-1-R | AGCTTCGTGGTGCATCTC | qPCR amplification of predicted EF-1α gene | XM_004240531.4 | qPCR reference gene | Roternberg et al., 2006 |
| UBI3-F | GCCGACTACAACATCCAGAAGG | qPCR amplification of *UBI3* | NM_001346406.1 | qPCR reference gene | Roternberg et al., 2007 |
| UBI3-R | TGCAACACAGCGAGCTTAACC | qPCR amplification of *UBI3* | NM_001346406.1 | qPCR reference gene | Roternberg et al., 2008 |
| GAME31-VIGS-F-XbaI | CGGTCTAGATGAGAGTTTGTGTATTCCTG | amplication of predicted *GAME31* amplicon for restriction enzyme cloning | XM_004233493.3 | | this study |
| GAME31-VIGS-R-BamHI | CGGGGATCCCAAAACAACGTAAGAATTTG | amplication of predicted *GAME31* amplicon for restriction enzyme cloning | XM_004233493.3 | | this study |
| GAME5-VIGS-F-XbaI | CGGTCTAGATTGGTAGTATCCTTCATTCAC | amplication of *GAME5* amplicon for restriction enzyme cloning | NM_001361347.1 | | this study |
| GAME5-VIGS-R-BamHI | CGGGGATCCGGTTCTGGAACTGTATTATTG | amplication of *GAME5* amplicon for restriction enzyme cloning | NM_001361347.1 | | this study |
| NOS-R | GATAATCATCGCAAGACCGGC | sequencing and PCR genotyping of VIGS-TRV2 constructs | n/a | | this study |
