## Supplemental Figures 1, 2, 3, 4, 5, 6, 7 for "Steroidal glycoalkaloids contribute to anthracnose resistance in *Solanum lycopersicum*"

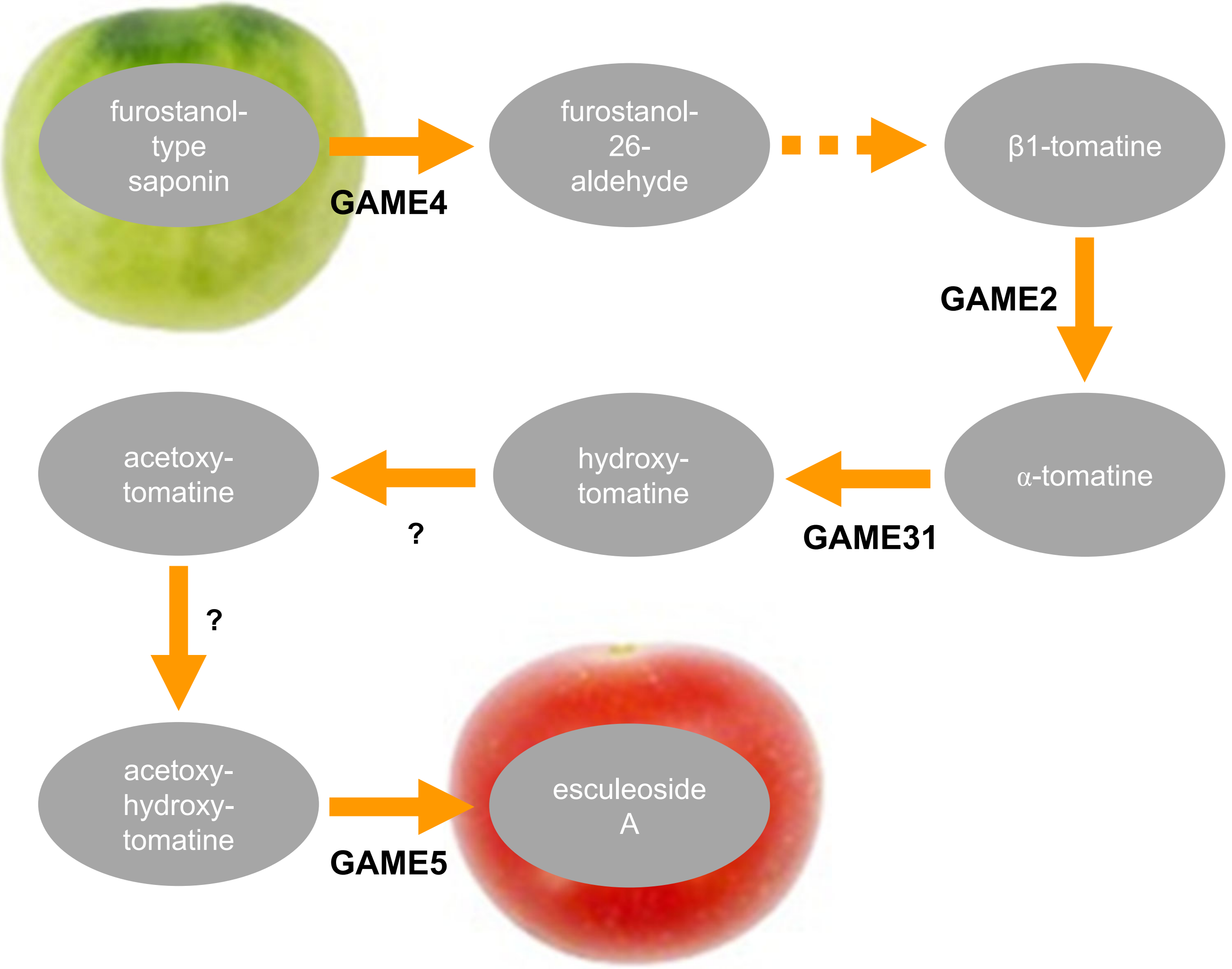


**Figure S1.** Simplified schematic of the steroidal glycoalkaloid (SGA) metabolic pathway during tomato fruit ripening. Biosynthetic reactions are catalyzed by glycoalkaloid metabolic enzyme (GAME) genes. Unripe tomato fruit is characterized by high levels of α–tomatine and other SGAs synthesized in earlier steps of the pathway, and red ripe fruit is characterized by high levels of the end product esculeosides.

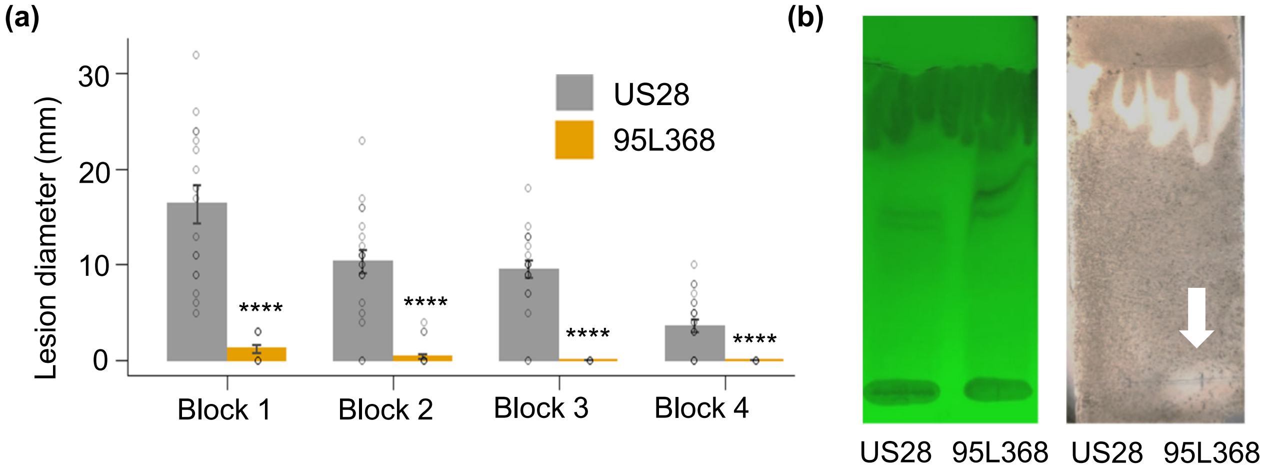


**Figure S2.** *Colleotrichum* resistance in the tomato accession 95L368. (a) Lesion diameter following *Colletotrichum* infection. Ripe fruit (n $\geq$ 13) from each block were harvested and inoculated with *Colletotrichum* and lesion size measured at 9 dpi. Data points represent individual biological replicates, and error bars indicate SEM. Asterisks indicate statistical significance ($p < 0.0001$, 1-way Student's T-test). (b) TLC bioassay. Reverse-phase separated, polar fractions of ripe fruit tissue were further separated via TLC (left), then inoculated with *Colletotrichum* and incubated for 3-4 days (right). The white arrow highlights a zone of inhibition observed for the 95L 368 sample.

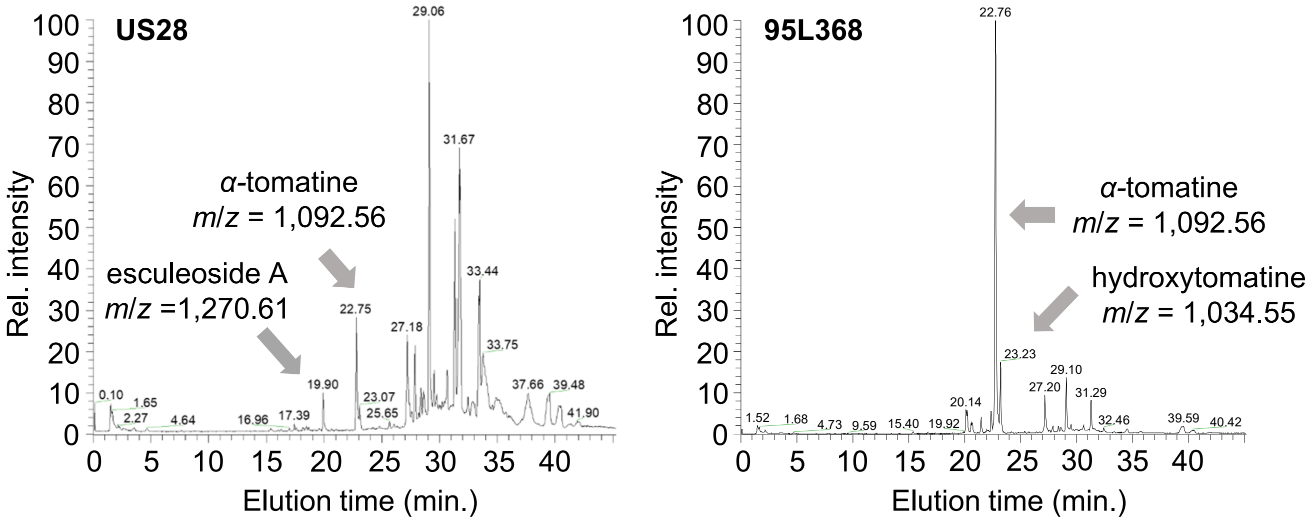


**Figure S3.** 95L368 accumulates enhanced levels of SGAs. Representative TICs from electrospray positive ionization HPLC-MS of ripe red fruit tissue samples of US28 (left) and 95L368 (right).

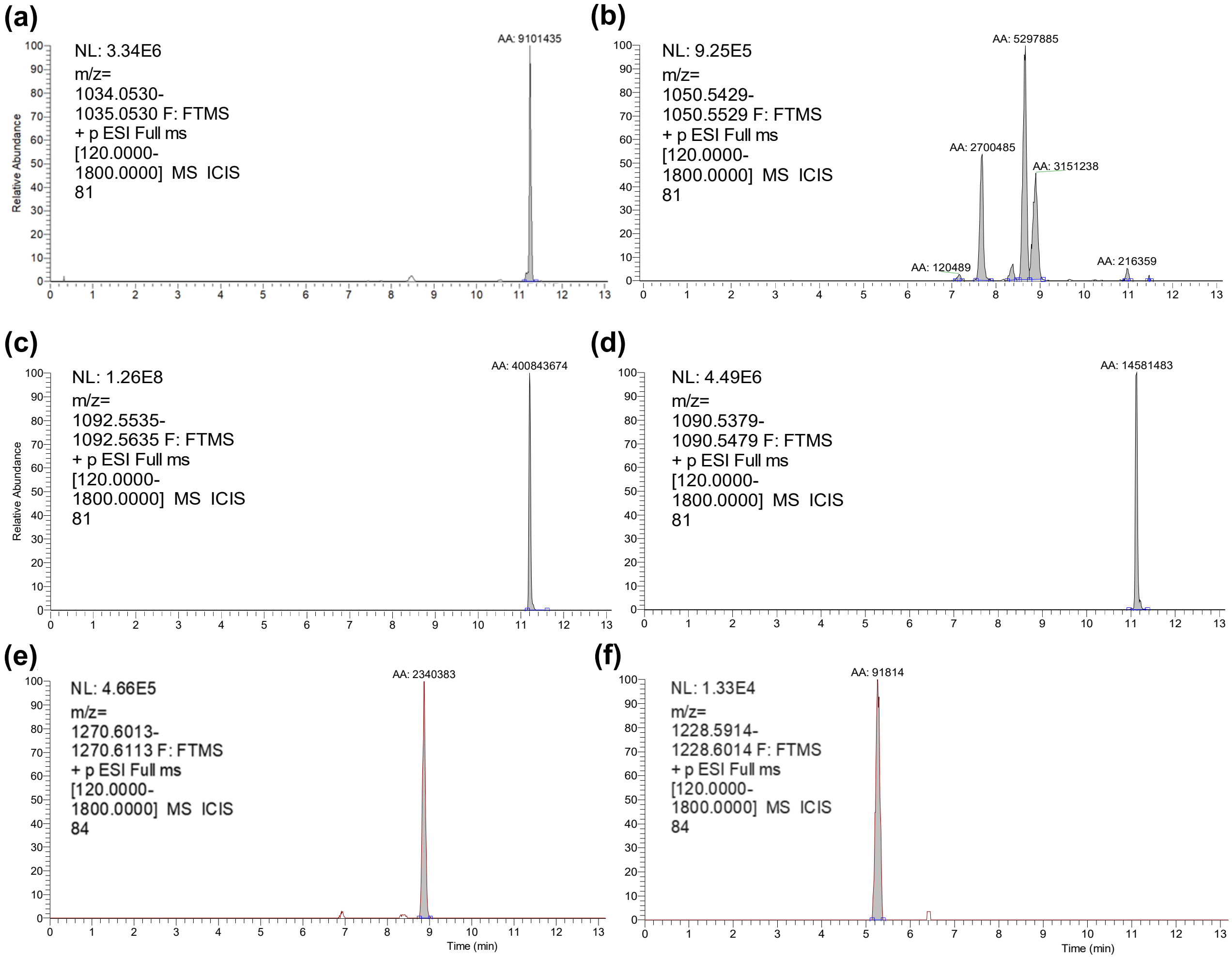


**Figure S4.** Representative extracted ion chromatograms (EICs) for detected SGAs. (a) α–tomatine. (b) Total hydroxytomatine isoforms. (c) Acetoxytomatine. (d) Dehydro-acetoxytomatine. (e) Esculeoside A. (f) Esculeoside B. Methanol extractions of lyophilized samples of red ripe fruit were separated and analyzed via UHPLC-HRMS, with masses calculated from α–tomatine standard. EICs (a) – (d) were generated from samples of 95L368. EICs (e) – (f) were generated from samples of US28.

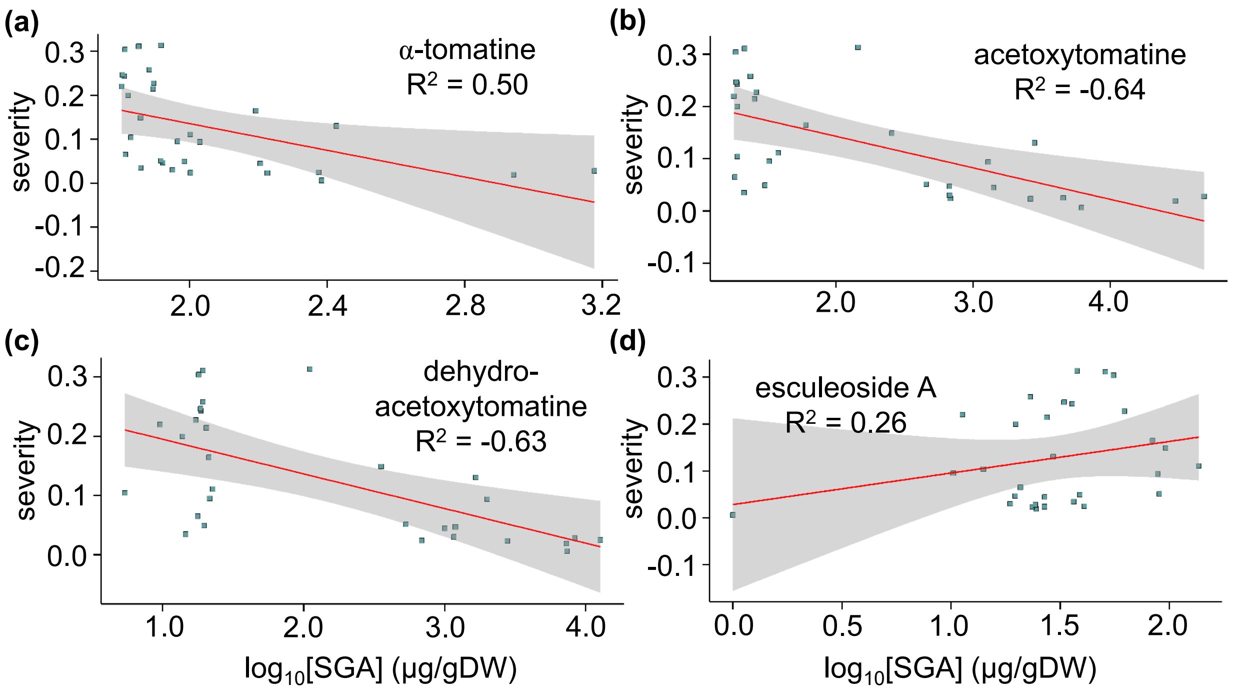


**Figure S5.** Mean detected SGAs vs. mean anthracnose disease severity. (a) α–tomatine. (b) Acetoxytomatine. (c) Dehydro-acetoxytomatine. (d) Esculeoside A. Following *Colletotrichum* infection, anthracnose lesion diameters were transformed to disease severities via normalization to the within-block maximum lesion size. Mean SGA levels were UHPLC-HRMS analysis of methanol extractions and $\log_{10}$-transformed. Each point corresponds to an individual parent line or RIL. Trendlines were generated via GLM and the grey regions represent the 95% C.I.

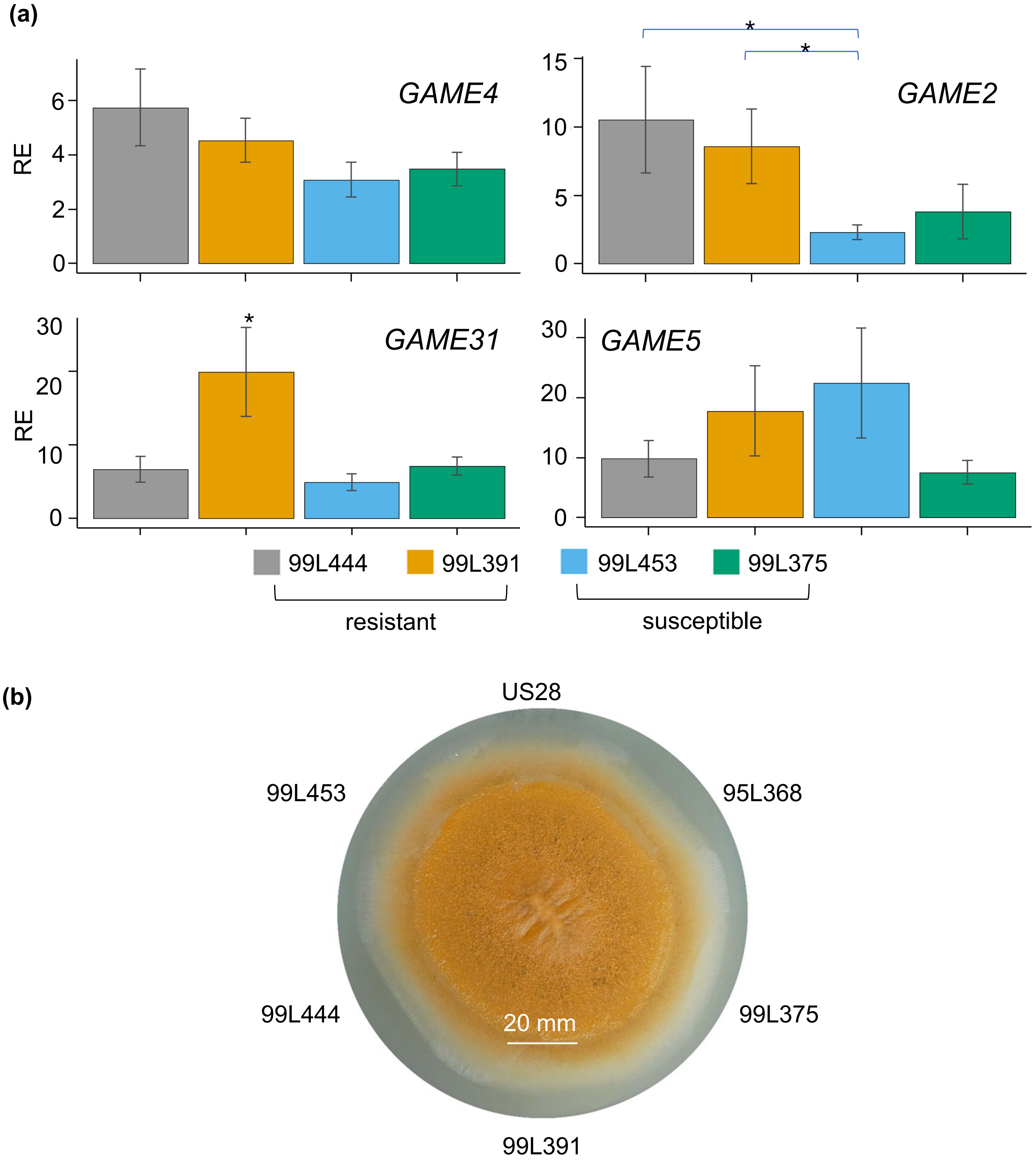


**Figure S6.** Anthracnose-resistant and –susceptible RILs exhibit variable GAME gene expression and *Colletotrichum* growth inhibition. (a) GAME gene expression. *GAME4* (upper left) and *GAME2* (upper right) expression was measured in mature green fruit tissue, and *GAME31* (lower left) and *GAME5* (lower right) expression was measured in ripe red fruit tissue. Asterisks indicate statistical significance ($p < 0.05$, 1-way Student's t-test) and error bars indicate SEM. For each target, qRT-PCR was conducted with 2 technical replicates of 3 biological replicates per line, in  independent experiments. (b) *Colletotrichum* plate growth inhibition assay. *Colletotrichum* strain C9 was inoculated onto V8 agar plates, with 80 µL red fruit tissue methanol extractions aliquoted to the perimeter of the plate at 7 dpi, and photographs taken at 5 dpt. This assay was repeated to similar results.

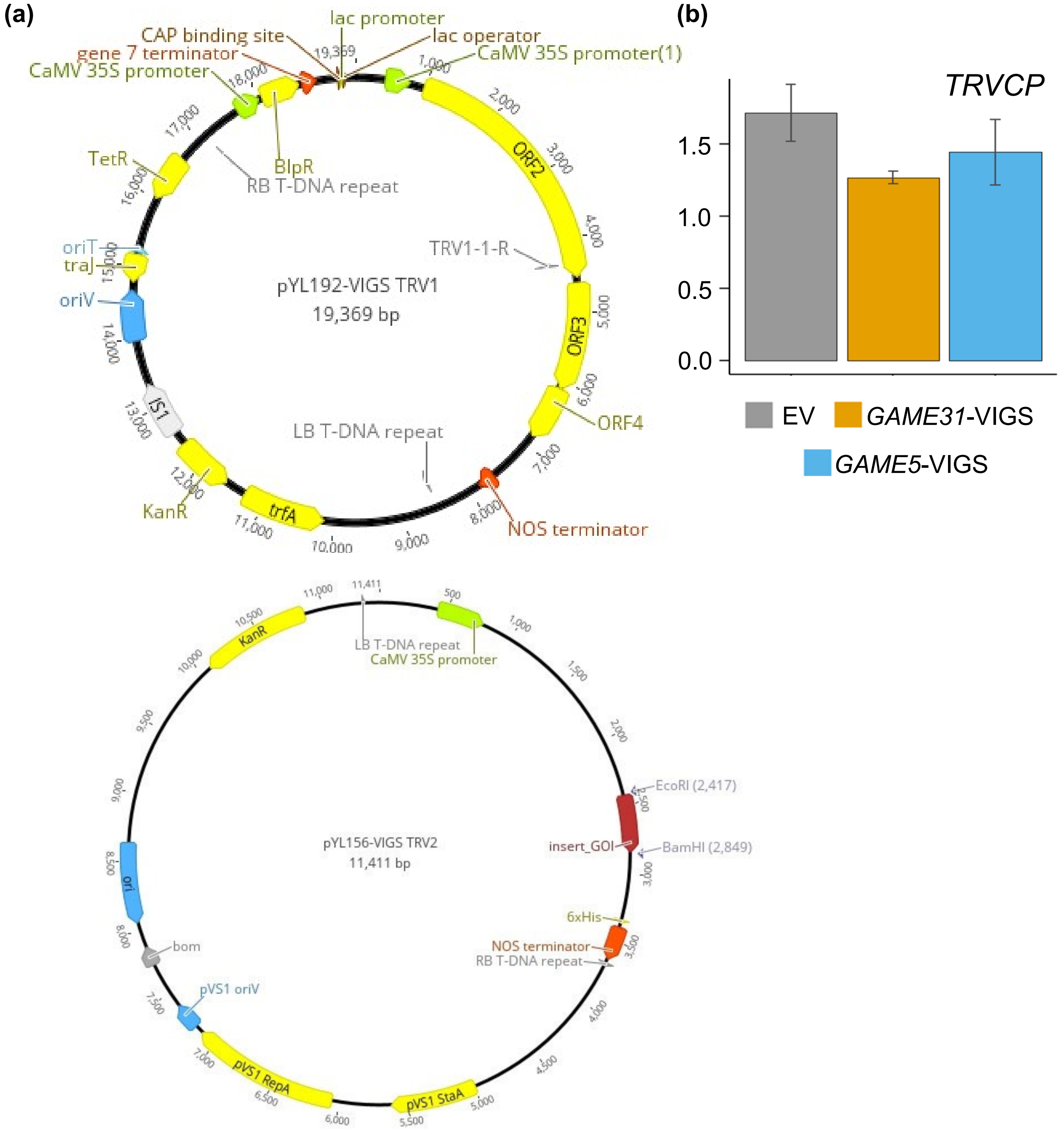


**Figure S7.** Transient silencing of *GAME* genes via VIGS. (a) TRV1 (upper) and TRV2 (lower) plasmids utilized in the bipartite VIGS system. (b) Validation of TRVCP expression in empty vector (EV), *GAME31*-VIGS, and *GAME5*-VIGS transformants. Samples from infected fruit were collected at 7 dpi and processed for RNA extraction. For each target, qRT-PCR was conducted with 2 technical replicates of 3 biological replicates per treatment. EV, empty vector control. Error bars indicate SEM.
